# Long-term Effects of Early Life Adversity On Brain Dopamine and Serotonin Receptor Systems Involved In Cocaine Reinforcement In Adult Macaques: a Positron Emission Tomography Study

**DOI:** 10.64898/2026.01.18.700208

**Authors:** Jose H. Acevedo-Polo, Erin R. Siebert, Jessica Khan, Mia I. Rough, Ronald J. Voll, Lahu N. Chavan, Mark M. Goodman, Jonathon A. Nye, Michael A. Nader, M. Mar Sanchez

## Abstract

Early life adverse (ELA) experiences such as child maltreatment (MALT) are associated with physical and mental illness, including substance use disorders (SUDs), but underlying neurobiological mechanisms remain unclear. This study examined long-term effects of infant MALT on adult brain serotonin (5HT) and dopamine (DA) receptors in corticolimbic regions involved in reward and emotional control, using positron emission tomography (PET) imaging, a translational infant MALT macaque model of cocaine use disorder (CUD) risk and a COC self-administration (SA) paradigm. The study focused on regional serotonin 5HT_1A_, 5HT_2A_, and dopamine D_2_/D_3_ receptor availability (BP) differences between MALT and Control animals using PET, both at baseline (pre-COC SA) and following chronic COC SA (once they reached a total of 100 mg/kg intake). We also examined whether levels of these neurochemical receptors predicted COC SA measures, including reinforcing effects and potency using fixed-ratio (FR) peak response rates and progressive-ratio (PR) peak breakpoint. Our findings showed long-term effects of infant MALT on 5HT, but not DA, receptors in corticolimbic circuits. Specifically, MALT animals showed lower 5HT_1A_ BP in the anterior cingulate cortex (ACC), medial prefrontal cortex (mPFC), and hippocampus compared to Controls. A MALT by Sex interaction effect was detected in 5HT_2A_ BP in the OFC, with lower levels in MALT than Control males, but not in females. In addition, upregulation of 5HT_1A_ and 5HT_2A_ receptors was detected following chronic COC SA in most PFC subregions, hippocampus, and NAcc, particularly in the Control group. These findings suggest long-term effects of ELA on adult 5HT, but not DA, receptors in corticolimbic regions involved in emotional and reward processes. We also found associations between PET baseline (pre-COC SA) receptor BP data and COC SA measures. In particular, a positive correlation between 5HT_1A_ receptor BP in caudate and peak FR Response Rates, whereas amygdala 5HT_1A_ receptor levels were positively correlated with peak PR breakpoint and negatively correlated with peak FR Response Rates. Overall, these findings suggest an important role of 5HT_1A_ and 5HT_2A_ PFC receptors in early COC-related changes in reward circuitry and of amygdala 5HT receptors on cocaine-maintained behaviors. The dynamic change of these 5HT_1A_ and 5HT_2A_ receptors following chronic COC exposure was blunted in animals with ELA. It would be important to understand the biological consequences of these dynamic changes in 5HT receptors and whether they are associated with other stages of the addiction cycle, for example COC relapse, which could inform future pharmacological interventions that target 5HT receptors for treatment of CUD.

**Simple Summary:** We studied the long-term effects of early life adversity (ELA) on adult brain dopamine (DA) and serotonin (5HT) signaling in corticolimbic regions involved in emotional and reward regulation. We used specific PET radioligands that bind to the DA D2/D3, 5HT_1A_ and 5HT_2A_ receptors, finding lower levels of 5HT, but not DA, receptors binding potential (BP) in animals that experienced ELA. We also found associations between PET receptor BP measures and reinforcing effects of cocaine in i.v. self-administration paradigms using fixed- and progressive-ratio reinforcement schedules. In addition, a strong upregulation of 5HT, but not DA, receptors was identified following chronic cocaine exposure in prefrontal cortex (PFC). Our findings suggest long-term effects of ELA on adult PFC 5HT_1A_ and 5HT_2A_ receptors. The findings also suggest an important role of 5HT_1A_ and 5HT_2A_, more so than D2/D3, receptors in early cocaine-related changes in reward circuitry. The early dynamic changes of these 5HT receptors could serve as biomarkers for cocaine use disorder (CUD) and inform future pharmacological interventions.

## Introduction

Early life adversity (ELA) is a major risk factor for physical and mental illness (Carr et al., 2013; Cicchetti & Toth, 2005; Douglas et al., 2010; Sinha, 2008). Childhood maltreatment (MALT) is a form of ELA that has been linked with the development of psychiatric disorders, such as anxiety, depression, post-traumatic stress disorder (PTSD), and substance use disorder (SUD) (Chu et al., 2013; Fergusson et al., 2008; Sanchez et al., 2001; Widom, 1999; Young & Tandon, 2025). More than 20 million Americans suffer from SUD, where overdose deaths represent only a fraction of their negative health consequences (Volkow & Boyle, 2018) and ELA is a known vulnerability factor, including for shorter abstinence length (Greenfield et al., 2002) and susceptibility to relapse (Heffner et al., 2011), as well as for reduced treatment-compliance (Jaycox et al., 2004) across different substance of abuse including alcohol, nicotine, opioids, and psychostimulants such as methamphetamine and cocaine -COC- (Enoch, 2011; Pilowsky et al., 2009; Sinha, 2008). Although decades of preclinical work support the association between ELA and SUDs, (Higley et al., 1991; Kirsch & Lippard, 2022; Kosten et al., 2000; Moffett et al., 2007; Vazquez et al., 2006; Wakeford et al., 2018), the underlying neurobiological mechanisms of increased vulnerability to psychostimulant addiction, such as COC, remain unclear. One possibility is that COC use disorder (CUD) is, at least in part, due to comorbid deficits in reward processes as well as in emotional and stress regulation caused by ELA-induced alterations in underlying neurocircuits. Specifically, a hyperactive stress-response system due to ELA may drive changes in reward-related circuitry resulting in a neurobiological system that is sensitive to the maladaptive reinforcing effects of addictive substances and to relapse (Wakeford et al., 2018). Previous studies showed that deficits in emotional regulation existed alongside stress-related disorders that were co-morbid with SUDs (Hyman et al., 2007; Simpson & Miller, 2002). This is supported by preclinical evidence in non-human primates (NHP) showing that ELA leads to elevated levels of the stress hormone cortisol and reduced brain serotonin (5HT) function/turnover, which are associated with excessive alcohol consumption in adulthood (Fahlke et al., 2000; Higley et al., 1991).

ELA can affect brain structural, functional and neurochemical development including alterations in brain 5HT and dopamine DA neurotransmission and their receptors, which are critical for reward, emotional and stress regulation (Teicher et al., 2003). ELA has been associated with disruptions in prefrontal cortex(PFC)-striatal dopaminergic reward circuits (Smith & Pollak, 2020). Infant MALT, in particular, can affect the development of the structure and connectivity of stress, emotional and reward regulation circuits, including the amygdala, PFC and nucleus accumbens (NAcc) across different species (Wakeford et al., 2018). These alterations include reduced PFC-amygdala connectivity in rhesus (Morin et al., 2020), and between the PFC, caudate, and putamen (Yan et al., 2017), as well as structural and functional alterations in the NAcc in rats (Monroy et al., 2010; Peña et al., 2014; Smith & Pollak, 2020).

The strong PFC connections to the NAcc are critical for reward and motivation (Camara et al., 2009), and their alterations contribute to increased risk of initiation of drug abuse following ELA (Balouek et al., 2023; Wakeford et al., 2018). The NAcc plays a major role in motivation, reward processing, the regulation of reward seeking and drug addiction (Everitt & Robbins, 2005; Koob & Volkow, 2010; Volkow et al., 2019; Xu et al., 2024). Prolonged stress and drug use cause maladaptive neuronal function in the NAcc circuitry which can eventually lead to pathological conditions (Xu et al., 2024). Alongside the PFC, the NAcc receives dopaminergic inputs from the ventral tegmental area (VTA) via the meso-corticolimbic pathway, a critical circuit in motivation, cognition, and addiction (Reynolds & Flores, 2021).

The PFC also plays a critical role in regulating stress and emotional reactivity through its connections with the amygdala, which plays a central role in threat detection, fear learning (Mah et al., 2016) and triggering the stress response (LeDoux, 1994; Rodrigues et al., 2009). Due to the amygdala bidirectional connections with the PFC that have been evolutionarily conserved (Gangopadhyay et al., 2021), disruptions to the circuitry may impair the PFC’s ability to exert top-down control over amygdala appropriate emotional responses (Sun et al., 2023), as well as to mediate synergistic interactions based on affective and reward-related information provided through interconnections with the NAcc that allow goal-directed behaviors (Gangopadhyay et al., 2021). Thus, dysregulation caused by ELA to this communication may heighten sensitivity and emotional reactivity, further reinforcing maladaptive behaviors that add to alterations in reward processing circuits.

PFC subregions have specific roles in stress/emotional responses and reward. The orbitofrontal cortex (OFC) is critical in goal-directed behavior, inhibitory control of behavior and affects cognitive function by predicting outcomes to ensure rational decisions are made (Hiser & Koenigs, 2018). In response to prolonged exposure to stress and cortisol, morphological and metaplastic changes in OFC neurons can occur, including reduction in dendritic spine density in basal dendrites of pyramidal neurons and in dendritic spine volumes and lengths (Barfield et al., 2020). The dorsolateral PFC (dlPFC) controls executive functions like working memory, value encoding, cognitive flexibility and decision making (Panikratova et al., 2020) and stress reduces dlPFC activity and functional connectivity (FC) in human fMRI studies (Joyce et al., 2024). The ventrolateral PFC (vlPFC) interfaces with sensory and motor areas while receiving emotional and motivational information from the OFC and amygdala (Barbas & Deolmos, 1990; Ilinsky et al., 1985). It plays a critical role in response inhibition, cognitive control through reappraisal and suppression of emotions and stress alters its FC with amygdala potentially influencing emotional processing and stress responses (Quaedflieg et al., 2015). The ventromedial PFC (vmPFC) is involved in adaptive stress responding and higher-order executive function such as decision making (Girotti et al., 2018; Maier & Watkins, 2010). fMRI studies have shown that increased vmPFC activity during stress was associated with active coping, whereas blunted vmPFC dynamic activity was associated to maladaptive coping, including binge drinking, emotional eating, irritability and fights (Sinha et al., 2016). Areas in the medial PFC (mPFC) -e.g. Brodmann Areas (BA) 10, 32-play an essential role in cognitive processes, emotional regulation, sociability and motivation by integrating information from limbic structures and other PFC regions, updating it and sending it back to output structures (Xu et al., 2019). Increases in mPFC activity and plasticity are critical for restoring adaptive coping mechanisms and executive function following chronic stress (Fucich et al., 2018). The subgenual cingulate cortex (SGC) -BA 25 in humans-is a separate component of the vmPFC and an important node in a network of corticolimbic regions (Hamani et al., 2011) that serves as the output from the frontal lobe to regions like the hypothalamus to regulate endocrine, visceral, appetitive and emotional states (Joyce et al., 2020; Öngür & Price, 2000), effectively controlling autonomic and somatic responses to stress (Alexander et al., 2020). The anterior cingulate cortex (ACC) is important for stress and emotional regulation, reward and motivation. It interconnects and integrates neurons from the frontal cortex, thalamus, and amygdala and also processes cognitive, emotional and autonomic functions (Yalcin et al., 2014). The ACC is associated to the pathophysiology of depression and pain processing (Yalcin et al., 2014).

Although DA and 5HT receptors are important for stress and emotional regulation, as well as for motivation and reward processes affected in psychiatric disorders (Akimova et al., 2009; Dunlop & Nemeroff, 2007; Perani et al., 2008; Stockmeier, 2003), there is less information available on the long-term impact of MALT and other ELA experiences on the brain DA and 5HT involved in those functions. The DA D_2_/D_3_ receptors are often inhibitory (Pivonello et al., 2007) and interesting due to their primary location in the NAcc, VTA, hippocampus, and frontal cortex (Ayano, 2016; Klein et al., 2019) and not just a major role in locomotor control (Plaisance et al., 2024), but also for addictive behaviors based on their strong role in reward, motivation and reinforcement (Plaisance et al., 2024). Chronic exposure to drugs of abuse leads to adaptive changes in D_1_, D_2_, and D_3_ receptor density, distribution and sensitivity, especially in the NAcc (Plaisance et al., 2024), particularly a downregulation in these DA receptors, resulting in reduced sensitivity to DA release and drug tolerance (Solinas et al., 2019; Volkow et al., 2019). Among the 7 types of 5HT receptors comprising a total of 14 subtypes (Sharp & Barnes, 2020), two are of particular interest for this study due to their roles at the interface of stress/emotional regulation and reward: 5HT_1A_ receptors (exhibit inhibitory control over firing and release of 5HT) and 5HT_2A_ receptors (exhibit excitatory effects over firing and release of 5HT), which are widely distributed and positioned to modulate DA transmission (Howell & Cunningham, 2015; Schenk & Highgate, 2021). These 5HT receptors are located across key reward circuitry brain regions, including the VTA (Doherty & Pickel, 2000), NAcc (Mengod et al., 1990), and PFC (Celada et al., 2013; Martín-Ruiz et al., 2001; Mengod et al., 1990; Pazos & Palacios, 1985). For example, in the PFC, 5HT_1A_ postsynaptic receptors are localized on pyramidal neurons where activation would hyperpolarize these neurons causing a decrease in firing activity (Schenk & Highgate, 2021), which can diminish glutamatergic output from the PFC to regions like the NAcc (Meunier et al., 2013), therefore decreasing reward processes and reinforcing effects mediated by DA release elicited by stimulants such as COC (Howell & Cunningham, 2015). On the other hand, 5HT_2A_ receptors are localized mainly on dendrites of PFC pyramidal output neurons (glutamatergic) and in GABAergic interneurons but activation of these receptors depolarizes these neurons, effectively increasing their excitability (Schenk & Highgate, 2021). PFC pyramidal neurons with 5HT_2A_ receptors project to both the NAcc and the VTA (Mocci et al., 2014; Vázquez-Borsetti et al., 2009); thus, activation of these receptors causes increased glutamatergic activation of NAcc and VTA, increasing the reinforcing effects of COC by increasing DA release in both these regions. Furthermore, 5HT_1A_ and 5HT_2A_ receptors are located on VTA DA neurons that project to the NAcc and PFC, further regulating DA synthesis and release.

Given the roles of 5HT_1A_ and 5HT_2A_ receptors in stress/emotional regulation and reward circuits and their modulation of DA signaling, alterations in these receptors have been implicated in psychiatric disorders, including anxiety, depression and SUD. For example, lower 5HT_1A_ receptor PET binding potentials (BP) are found in limbic and cortical regions in depressed patients, while 5HT_2A_ receptor (Savitz & Drevets, 2013) and D_2_/D_3_ receptors BP results are inconsistent (Savitz & Drevets, 2013). However, despite growing evidence linking ELA to neurodevelopmental alterations of stress/emotional regulation and reward circuits, the relationship between ELS-induced monoaminergic dysregulation, specifically in the context of MALT, and the role of 5HT receptors modulating DA signaling in CUD remains understudied. COC binds to dopamine transporters (DATs), acting as a reuptake inhibitor of DA into the presynaptic terminal (Rasmussen et al., 2001; Verma, 2015), resulting in an increase of DA in the synaptic cleft that leads to enhanced stimulation of postsynaptic receptors and subsequent reinforcing effects. COC also binds to 5HT transporters (SERT) and norepinephrine transporters (NET), boosting synaptic levels of 5HT and NE respectively by inhibiting their reuptake. Due to the different ways that COC interacts with different neurotransmitter systems, there are other mechanisms apart from DA modulation that contribute to the behavioral and reinforcing effects of COC, including intricate relationships between 5HT and DA in drug reinforcement. Further research exploring these interactions could inform the development of new pharmacotherapies for CUD.

To understand the neurobiological mechanisms underlying ELA increased risk for SUDs it is important to use prospective, longitudinal studies, but this is very challenging in children at-risk and there are many confounding factors (e.g. SES, prenatal exposure to drugs and/or stress) at play. Using rhesus macaques as a model of ELA and drug abuse is a powerful translational strategy to understand how ELA alters the structural and functional development of reward and emotion regulatory circuits, increasing the risk of drug abuse. Additional strengths of using rhesus macaques as model of MALT include the ability to apply brain magnetic resonance imaging (MRI) and PET techniques, the fact that neurodevelopmental processes are around four times faster than in humans, and the complex social behavior, including mother-infant relationships in this species, which translates to early human experiences (Howell & Sanchez, 2011; Sanchez, 2006; Sanchez et al., 2001). Therefore, the goal of this study was to examine the long-term effects of ELA due to MALT on increased risk for CUD using a COC self-administration (SA) paradigm in adult rhesus monkeys. The aim was to understand (1) the effects of MALT on brain 5HT_1A_ and 5HT_2A_ receptors and DA D_2_/D_3_ receptors in corticolimbic regions involved in reward and stress/emotional regulation using PET imaging; and (2) whether those neurochemical alterations were associated with COC SA measures.

## Methods

### Subjects, Housing and Infant Maltreatment Model

The study examined 22 adult rhesus monkeys (*Macaca mulatta*), ages 11-14 years. Animals were born and socially-housed at the Emory National Primate Research Center (ENPRC) Field Station, where half of them experienced early life stress (ELA) in the form of infant maltreatment (MALT) throughout the first 3 to 6 months of life (MALT: n = 13; 7 males, 6 females) and the other half were raised by competent mothers (Controls: n=9; 5 males, 4 females). These animals were part of a larger longitudinal study throughout infancy, juvenile, adolescent, and adult periods to examine the effects of infant MALT on brain development, behavior, emotional/stress regulation, DNA methylation, as well as risk for COC initiation and maintenance in adolescence and adulthood (Allen et al., 2025; Bronk et al., 2024; Drury et al., 2017; Howell et al., 2019; Lardenoije et al., 2025; McCormack et al., 2025; McCormack et al., 2022; Morin et al., 2020; Morin et al., 2019; Morin et al., 2025; Wakeford et al., 2020; Wakeford et al., 2019; Wakeford et al., 2024). The experimental timeline is outlined in **Suppl. Figure 1**. All animals lived with their mothers and families in complex social groups consisting of 75–150 adult females along with their juvenile and sub-adult offspring, as well as 2–3 adult males. These groups were housed in outdoor compounds with access to indoor, climate-controlled, areas and were fed standard high fiber, low fat monkey chow (Purina Mills Int., Lab Diets, St. Louis, MO) in addition to seasonal fruits and vegetables provided twice daily alongside enrichment items. Water was available *ad libitum* and enrichment items were provided as part of standard care. Infant MALT was operationalized as the co-morbid experience of physical abuse (violent behaviors such as dragging, crushing, throwing or stepping on the infant) and rejection of the infant by the mother during the first three months of life. These are aberrant and negative maternal behaviors that cause pain, behavioral distress, and elevations of the stress hormone cortisol in rhesus infants (Drury et al., 2017; Howell et al., 2013; Maestripieri, 1998; Maestripieri & Carroll, 1998; Maestripieri, Higley, et al., 2006; Maestripieri, McCormack, et al., 2006; Maestripieri et al., 2000; McCormack et al., 2009; McCormack et al., 2006; McCormack et al., 2022; Sanchez, 2006) and are never exhibited by Control mothers, who showed competent care (e.g. cradling, ventral contact, protection of the infant (McCormack et al., 2015). To disentangle the effects of maternal care from potential confounding heritable factors of MALT phenotypes, a cross-fostering experimental design was used to randomly assign infants at birth to rearing group (i.e. to foster mothers with histories of infant MALT or of nurturing maternal care -Controls-; (Drury et al., 2017; Howell et al., 2017; McCormack et al., 2022). **Table 1** provides details on the cross-fostering groups in this study. Mother-infant pairs were chosen from different social ranks, matrilines, and fathers to provide genetic and social rank diversity in the various groups. Following cross-fostering, maternal care data were collected during the first three postnatal months to confirm and quantify abuse and rejection rates, as well as other maternal behaviors, mother-infant interactions and infant’s behaviors (see McCormack et al., 2022 for details on behavioral data collection and analyses). Focal observations (4×30min) were collected by trained coders (>90% interrater reliability) at different infant ages, on separate days (5 days/week during month 1; 2 days/week during month 2; 1 day/week during month 3), totaling 16 hours mother-infant relationship data per pair across the first 3 postnatal months. Observations were collected in real time using a modified rhesus monkey ethogram (Altmann, 1962; McCormack et al., 2006) and recorded via *WinObs*, an in-house behavioral coding software developed at the ENPRC (Graves & Wallen, 2006). These behavioral methods were selected and optimized for documenting early infant MALT, as physical abuse is the highest during the first postnatal month and stops by the third month, and high levels of infant rejection are exhibited by MALT mothers during the first 3 months in comparison to Controls (McCormack et al, 2022).

**Table 1.**
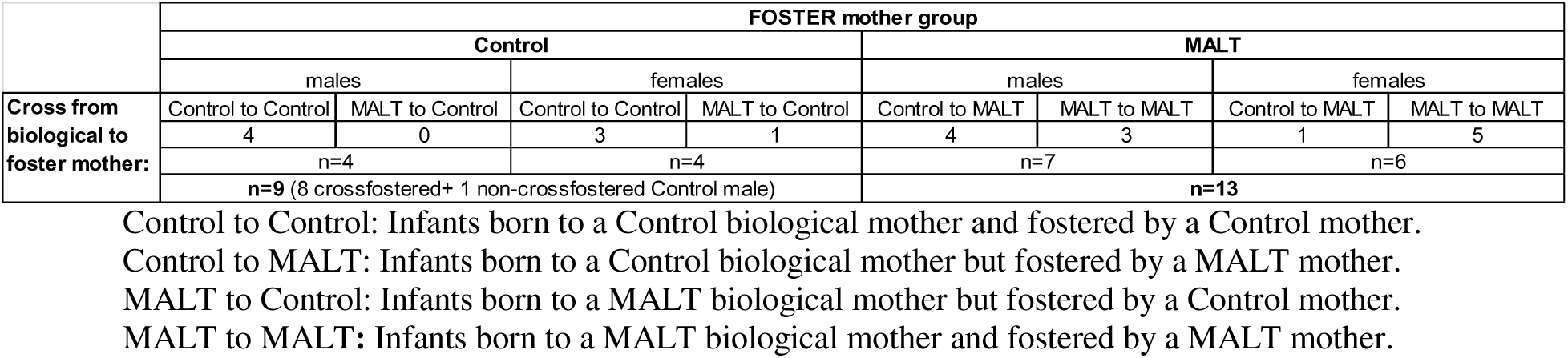
Sex Distribution and Group Assignment Based on Randomized Cross-Fostering at Birth to a Control or Maltreating (MALT) Foster Mother and Biological Mother.

At 4-5 years of age, the 22 animals in this study were transferred to the ENPRC Main Station where they were studied during adolescence and adulthood. The animals were either pair- or single-housed (due to incompatibility/aggression with partner), counterbalancing for Control and MALT group, and maintained under controlled temperature (22 ± 2°C, humidity: 25–50%) with a 12-hour light/dark cycle (lights on at 0700 and off at 1900). They were fed Purina monkey chow (Ralston Purina, St. Louis, MO, USA), supplemented daily with vegetables and fruits and water available *ad libitum*. Environmental enrichment was provided regularly. After several months of acclimation, animals underwent behavioral (baseline startle)/cognitive tasks, neuroendocrine assessments, brain magnetic resonance imaging (MRI) and positron emission tomography (PET) scans before COC SA studies during adolescence. Following approximately 3-4 years of abstinence the startle, neuroendocrine, brain MRI/PET and COC SA studies were repeated in adulthood. All study protocols and animal care complied with the National Institutes of Health Guide for the Care and Use of Laboratory Animals (8^th^ edition, revised 2011), followed recommendations from AAALAC International, and were approved by the Emory University Institutional Animal Care and Use Committee (IACUC). The ENPRC is fully accredited by AAALAC International.

### Brain Positron Emission Tomography (PET) Data Acquisition, Processing and Analysis

PET scans were acquired during adulthood with a microPET Focus 220 scanner (Concord Microsystems, Knoxville, TN), featuring a 8/26cm axial/tranaxial field-of-view and an isotropic reconstructed resolution of 1.7 mm in all directions. These PET scans were acquired at (1) baseline (pre-COC SA) in all 22 animals, and (2) again right after completion of adult COC SA studies, once each subject had reached a cumulative COC intake of ≥100mg/kg (post-COC SA PET scan); see **Suppl. Figure 2** for schematic diagram showing the timeline of these studies. Only 13 out of the 22 animals completed the COC SA fixed-ratio (FR) schedule studies and 11 out of those completed the progressive-ratio (PR) studies, as published elsewhere (Allen et al, 2025). Post-COC SA PET scans were collected in 9 of those 11 animals by the time this study was completed. Animals were anesthetized with a telazol/ketamine cocktail, followed by endotracheal intubation and maintained under anesthesia with isoflurane (1-2%, inhalation) during the PET scans. The animals were scanned supine and transmission data were collected with a Co-57 point source for purposes of attenuation correction of the emission data. PET scans were performed at 1-2 week intervals with the following PET ligands injected intravenously (IV) in counterbalanced order: [18F]fallypride for the D_2_/D_3_ receptor (Mukherjee et al., 1999), [18F]MPPF for the 5HT_1A_ receptor (Le Bars et al., 1998), and [18F]MDL 100,907 for the 5HT_2A_ receptor (Chavan et al., 2023). PET data were acquired for 120 minutes for each PET ligand and divided into 21 time-series frames for kinetic analysis. PET data was reconstructed using a 3D ordered subset expectation maximization and maximum a posteriori regularization algorithm (Yang et al., 2004) to a matrix size of 128 x 128 x 990 and voxel size of 0.9 mm in all directions.

### PET Data Analysis

PET data and analyses were done following published protocols by our group for rhesus brain (Wakeford et al., 2024). PET scans were corrected for intra-frame motion using an affine registration technique (Woods et al., 1992) and co-registered to the individual subject T1-weighted MRI scan by optimizing the mutual information metric (Maes et al., 1997). Individual MRI scans were spatially normalized to the UNC-Emory 12-month rhesus T1-MRI brain atlas (Shi et al., 2017) using SPM8 software with parameter settings optimized for rhesus (McLaren et al., 2010). Brain regions of interest (ROIs) defined in the UNC-Emory 12-month atlas were transformed to each animal’s native T1 MRI space using the inverse non-linear transformation matrix generated from SPM8 Segment method. Time-activity curves (TACs) were derived for each ROI. Subsequent analyses were conducted using an in-house pipeline developed by our group in the International Data Language -IDL- environment (Harris Geospatial Solutions Inc., Broomfield, CO). Binding potential (BP) estimates (i.e. non-displaceable BP: BPND) were defined as a ratio of specifically bound radioligand to the concentration of non-displaceable radioligand in tissue (Turkheimer et al., 2012) and calculated using established kinetic modeling techniques where the cerebellum serves as the reference tissue input function (Embree et al., 2013; Michopoulos et al., 2013; Michopoulos et al., 2014; Wakeford et al., 2024). BPND of [18F]fallypride signal was determined via the multilinear reference tissue model (Christian et al., 2009). BPND of [18F]MPPF and [18F]MDL 100,907 signals were calculated using the simplified reference tissue model (Lammertsma et al., 1996). Although this PET study is based on BPND measures, from this point forward we will use “BP” as the main abbreviation for simplicity purposes. The following corticolimbic ROIs were included in the study because of their role in stress/emotional and reward circuits: amygdala, hippocampus, dorsal striatum (caudate, caudate), ventral striatum (NAcc), and PFC subregions: ACC, SGC, mPFC, vmPFC, OFC, dlPFC, and vlPFC. See **Suppl. Figure 3** for ROI anatomical criteria and examples of the label maps. Each ROI bilateral BP was calculated by taking the average of the left and right ROI BP.

### Cocaine Self-Administration (COC SA)

COC SA data used for association analyses are based in published findings (Allen et al., 2025) and summarized in the Results. Briefly, in *adolescence*, subjects were surgically implanted with IV catheters and underwent operant conditioning to self-administer COC hydrochloride dissolved in 0.9% sterile saline through the IV catheter. Data were collected on different phases of COC SA including acquisition, maintenance of drug intake, dose response curves (DRCs) under a fixed-ratio (FR) schedule of reinforcement (FR20), limited access, extended access, extinction, and reinstatement of COC SA response (Wakeford et al, 2019 and 2020). Although adolescent total COC intake was not significantly different between Control and MALT groups, it was included as a covariate in the statistical models in this study. Following a period of extended abstinence (approx. 3 years), in *adulthood* (Allen et al., 2025) the animals underwent COC SA again under FR20 to generate response rates (FR RR: responses/second) at different doses of the DRC (saline, 0.01, 0.003, 0.01, 0.03, 0.1, 0.3 mg/kg/injection). The COC dose associated with peak RR for each individual subject was one of the primary COC outcome measures in this study. Following determination of the COC DRC, the subjects transitioned to a progressive-ratio (PR) schedule to examine the reinforcing strength of COC. In this PR schedule, the number of responses (lever presses) necessary for reinforcement was systematically increased until the animal no longer received a reinforcer within a certain time (limited hold: LH 60min) or the 4hr session expired. The response requirement for each injection increases according to a published equation (Richardson & Roberts, 1996). Another COC outcome in this study was peak COC break-point or the maximum number of COC injections received at the peak of the DRC (saline, 0.01, 0.003, 0.01, 0.03, 0.1, 0.3 mg/kg/injection).

### Statistical Analysis

Analyses were conducted using IBM SPSS Statistics v.29.0 (IBM Corporation, Armonk, NY), with statistical significance set at p<0.05 for all tests. Variables were summarized as mean(x□)±standard error of the mean(SEM) in figures, and effect sizes were calculated as η_p_^2^. For statistical models assuming normality -all except for Spearman Rank’s correlations-, data were first checked for normality using the Shapiro-Wilk test and, if failed, data were log10-transformed for use in the statistical models. For clarity purposes we are only providing details on statistically significant findings in the Results section.

A Two-Way ANOVA was used to examine main and interaction effects of group (MALT, Control) and sex (male, female) on PET Baseline (pre-COC SA) 5HT_1A_, 5HT_2A_ and D_2_ /D_3_, receptor BP in each ROI. Post-hoc Bonferroni-corrected pairwise comparisons of the means were used when significant interaction effects were detected. For the group of animals that underwent COC SA, a repeated measures (RM) ANOVA was used to examine changes in ROIs 5HT_1A_, 5HT_2A_, and D_2_ /D_3_ receptors BP from Baseline (pre-COC SA) to post-COC SA with group (MALT, Control) as fixed factor and time (Baseline, post-COC SA) as the RM; sex was not included in these RM ANOVA models due to lack of power to detect interactions. Post-hoc Bonferroni-corrected pairwise comparisons of the means were used when significant interaction effects were detected. Although total COC intake during adolescence was not significantly different between Control and MALT groups it was included as a covariate in the statistical models to control for its potential confounding effects on the receptors BP.

For data reduction purposes for correlational analyses that examined associations between Baseline (pre-COC SA) 5HT_1A_, 5HT_2A_, and D_2_ /D_3_ receptor BP in the ROIs and COC SA outcome data (FR peak RR, PR Peak Breakpoint) a correlation-based clustering method was first used to identify clusters of associations between ROIs BPs for each PET ligand, using Spearman Rank’s correlation followed by visualization of the significant clustering results in correlation matrices (see **Suppl. Tables 1-3**). Clusters of strongly correlated ROIs BPs varied by ligand and we created composite scores (averages) of PET BP in ROIs included in each of the following clusters: (1) [18F]MPPF: Cluster 1 (dlPFC, mPFC, vmPFC) and Cluster 2 (ACC, OFC, SGC, mPFC), in addition to ROIs that did not cluster with others (Amygdala, Hippocampus, NAcc, Caudate, Putamen, vlPFC); (2) [18F]MDL100: Cluster 1 (ACC, dlPFC, mPFC, vmPFC, vlPFC), Cluster 2 (ACC, OFC), Cluster 3 (Amygdala, OFC, Caudate, Putamen), Cluster 4 (Amygdala, NAcc, OFC, SGC), and Cluster 5 (Hippocampus, ACC, Caudate, dlPFC, mPFC, vlPFC, vmPFC); (3) [18F]fallypride: Cluster 1 (ACC, OFC, SGC, mPFC, dlPFC, vlPFC, vmPFC) and ROIs that did not cluster with others (Amygdala, Hippocampus, NAcc, Caudate, Putamen). These ROIs BP clusters were then entered into Spearman Rank’s correlations to generate pairwise correlation matrices of associations between Baseline PET ligand clustered data and COC SA outcomes (FR peak Response Rate, PR peak Breakpoint).

## Results

### Baseline PET Binding Potential (BP) Analysis

Data were normally distributed except for the 5HT_1A_ receptor BP in Amygdala, Caudate, NAcc, Hippocampus, Putamen, and SGC, and D_2_/D_3_ BP in ACC, Amygdala, Caudate, Hippocampus, NAcc, Putamen, dlPFC, mPFC, and vlPFC, even following log-transformation. Homogeneity of variance was met for all data except for 5HT_1A_ receptor BP in Caudate, and 5HT_2A_ receptor BP in OFC, which failed even following log-transformation. Accordingly, analyses for these ROIs were conducted using log-transformed BP values to address deviations from normality and heteroscedasticity.

#### 5HT1A Receptor *BP*

Two-Way ANOVA revealed significant main effects of group (ELA) on 5HT_1A_ receptor BP in ACC (F_(1,18)_=5.159, *p*=.036, η_p_^2^=.223), Hippocampus (F_(1,18)_= 4.649, *p*=.045, η_p_^2^=.205), and mPFC (F_(1,18)_=6.132, *p*=.023, η_p_^2^=.254), with MALT animals showing significantly lower 5HT_1A_ receptor BP than Controls in these ROIs (**Figure 1**).

**Figure 1.**
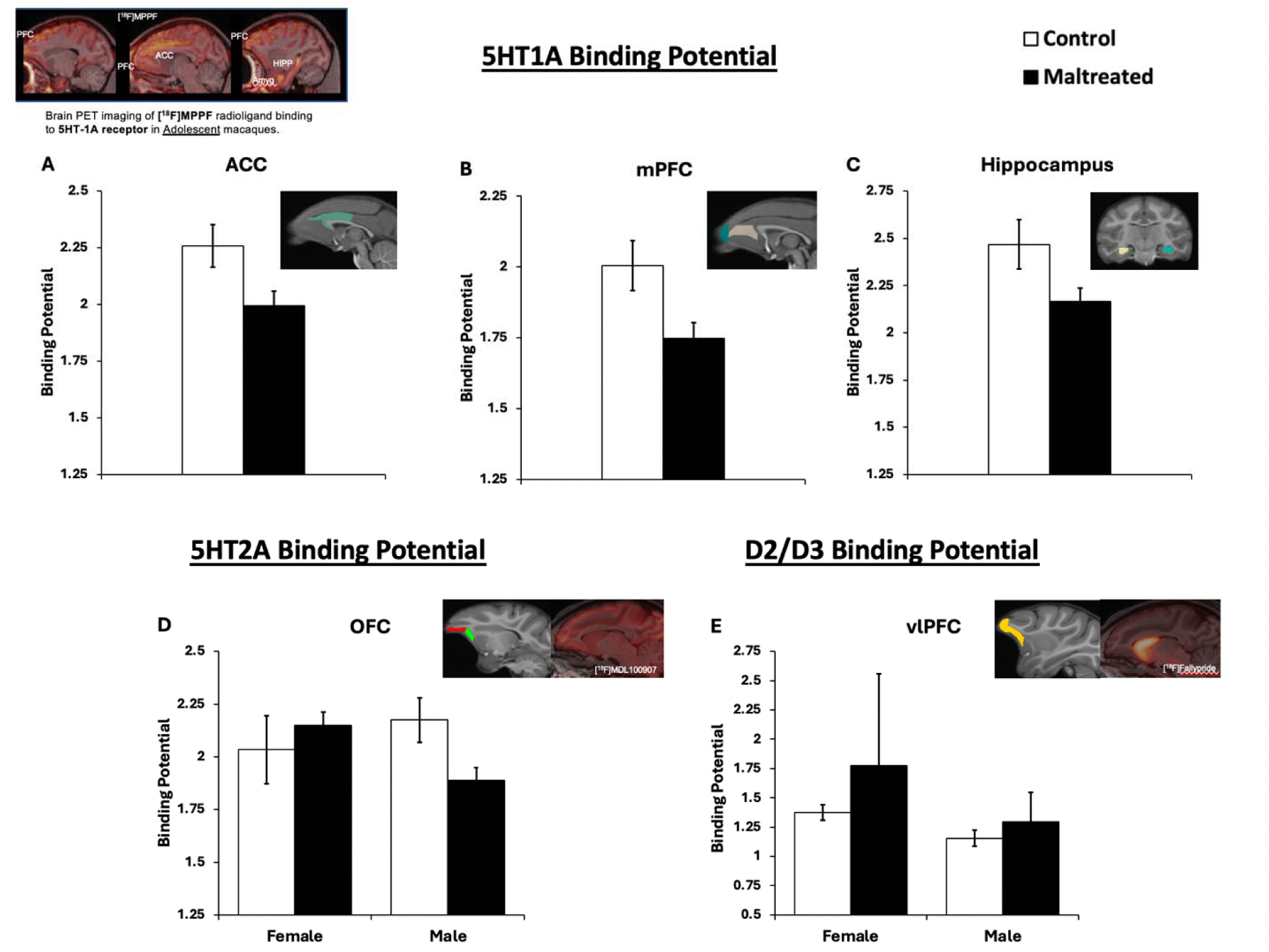
Baseline 5HT_1A_, 5HT_2A_, and D2/D3 Binding Potential (BP) in Anterior Cingulate Cortex (ACC), Medial Prefrontal Cortex (mPFC), Hippocampus, Orbitofrontal Cortex (OFC), and ventrolateral Prefrontal Cortex (vlPFC). Lower levels of 5HT_1A_ receptor were detected in Maltreated than Control adult rhesus monkeys in (A) ACC (F(1,18)=5.159, p=.036, ηp2 =0.327), (B) mPFC (F(1,18)=6.132, p=.023, ηp2 =0.254), and (C) Hippocampus F(1,18)=6.132, p=.023, ηp2 =0.254). A significant group x sex interaction effect was detected in (D) OFC (F(1,18)=5.013, p=.038, ηp2 =0.218), with lower BP in Maltreated than Control males (p<0.05, Bonferroni corrected post-hoc comparisons of the means), but not in females. A significant main effect of sex was found in (E) vlPFC (F(1,18)=5.052, p=.037, ηp2 =.219), with lower D2/D3 BP in males than females. Data represented as x□±SEM.

When the model was re-run with adolescence COC intake as a covariate in a Two Way ANCOVA, the main group effects remained in ACC (F_(1,17)_=5.175, *p*=.036, η_p_^2^=.223) and mPFC (F_(1,17)_=6.131, *p*=.024, η_p_^2^=.265), but disappeared in Hippocampus although with a trend towards significance (F_(1,17)_=4.291, *p*=.054, η_p_^2^=.202). In all these ROIs, MALT animals showed significantly lower 5HT1A Receptor BP than Control animals.

#### 5HT2A Receptor *BP*

Two-Way ANOVA revealed significant group by sex interaction effects on 5HT_2A_ receptor BP in the OFC (F_(1,18)_=5.013, *p*=.038, η_p_^2^=.218). Post-hoc tests showed significantly lower 5HT_2A_ receptor BP in MALT than Control males (p<0.05), while no differences were found for females (**Figure 1**).

The group by sex interaction effects disappeared in the OFC after adding adolescence COC intake as a covariate, although with a trend towards significance (F_(1,17)_=3.785, *p*=.068, η_p_^2^=.182). Adolescence COC intake had a significant effect as covariate in NAcc (F_(1,17)_=5.187, *p*=.036, η_p_^2^=.234), OFC (F_(1,17)_=5.088, *p*=.038, η_p_^2^=.230), SGC (F_(1,17)_=4.824, *p*=.042, η_p_^2^=.221), and vmPFC (F_(1,17)_=5.457, *p*=.032, η_p_^2^=.243).

#### D2/D3 Receptor BP

Two-Way ANOVA showed a significant main sex effect on D_2_/D_3_ receptor BP in the vlPFC (F_(1,20)_=5.052, *p*=.037; η_p_^2^=.219), with males showing lower BP than females (**Figure 1**). No other main or interaction effects were detected.

After adding adolescence COC intake as a covariate in the Two Way ANCOVAs the main sex effects remained in vlPFC (F_(1,17)_=5.802, *p*=.028, η_p_^2^=.254) with males showing lower BP than females.

### Post-cocaine self-administration PET BP Analysis

All data were normally distributed except for post-COC 5HT_2A_ receptor BP in ACC and D_2_/D_3_ receptor BP in Putamen, which failed normality even after log-transformation. Accordingly, analyses for these ROIs were conducted using log-transformed data. Homogeneity of variance was met for all data except for 5HT_2A_ receptor BP in Caudate, and D_2_/D_3_ receptor BP in ACC, Putamen, and dlPFC, which were run log-transformed.

#### 5HT1A Receptor *BP*

RM ANOVA revealed significant main effects of group in ACC (F_(1,7)_= 9.482, *p*=.018, η_p_^2^=.575), NAcc (F_(1,7)_=5.593, *p*=.050, η_p_^2^=.444), OFC (F_(1,7)_=12.234, *p*=.010, η_p_^2^=.636), SGC (F_(1,7)_=8.332, *p*=.023, η_p_^2^=.543), dlPFC (F_(1,7)_=19.426, *p*=.003, η_p_^2^=.735), and vmPFC (F_(1,7)_=15.886, *p*=.005, η_p_^2^=.694). In all these ROIs, MALT animals showed lower 5HT_1A_ receptor BP than Controls (**Figure 2**). Additionally, significant main time effects were found in Hippocampus (F_(1,7)_= 13.877, *p*=.007, η_p_^2^=.665), OFC (F_(1,7)_=33.017, *p*=.001, η_p_^2^=.825), SGC (F_(1,7)_=3.715, *p*=.001, η_p_^2^=.814), dlPFC (F_(1,7)_=85.745, *p*<.001, η_p_^2^=.575), mPFC (F_(1,7)_=25.920, *p*=.001, η_p_^2^=.787), vlPFC (F_(1,7)_=41.841, *p*<.001, η_p_^2^=.857), and vmPFC (F_(1,7)_=19.569, *p*=.003, η_p_^2^=.737). In all these ROIs, animals showed higher 5HT_1A_ receptor BP post-COC SA than at baseline (**Figure 2**). Significant group x time interaction effects were found in OFC (F_(1,7)_=41.849, *p*<.001, η_p_^2^=.857), SGC (F_(1,7)_=26.424, *p*=.001, η_p_^2^=.791), dlPFC (F_(1,7)_=13.818, *p*=.007, η_p_^2^=.664), and vlPFC (F_(1,7)_=7.056, *p*=.033, η_p_^2^=.602). Post-hoc tests showed that in the OFC and SGC, Control animals had significantly higher 5HT_1A_ receptor BP post-COC SA than at baseline (p<0.05) while no differences were found for MALT animals (**Figure 2**). On the other hand, in the dlPFC and vlPFC, both Control and MALT groups showed significantly higher 5HT_1A_ receptor BP post-COC SA than at baseline (p<0.05; **Figure 2)**. No other main or interaction effects were detected.

**Figure 2.**
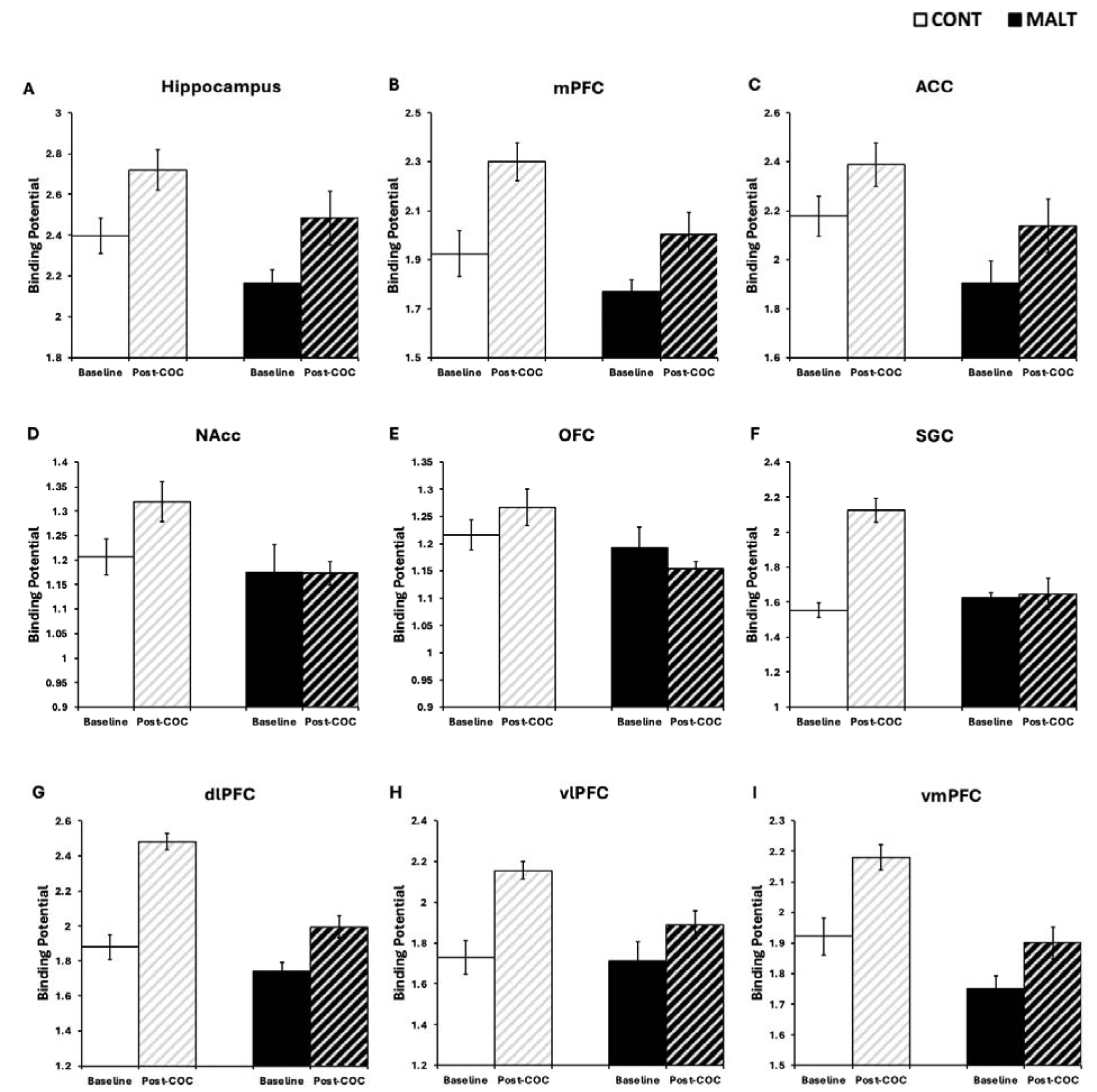
Changes in 5HT_1A_ receptor BP from Baseline (pre-COC SA) to post-COC SA in corticolimbic regions studied using [F18]-MPPF. Significantly main group effects were detected in NAcc, ACC, OFC, SGC, dlPFC, and vmPFC, with MALT animals showing lower receptor levels than Controls. Main significant time effects were also found in mPFC, vmPFC, OFC, SGC, dlPFC, vlPFC, with overall main receptor upregulation following COC SA. Group x time interaction effects were also detected in OFC, SGC, dlPFC, vlPFC. See statistical details in text. NAcc: Nucleus Accumbens ACC: anterior cingulate cortex, mPFC: medial prefrontal cortex; OFC: orbitofrontal Cortex, SGC: subgenual cingulate cortex, dlPFC: dorsolateral PFC, vlPFC: ventrolateral PFC, vmPFC: ventromedial PFC. Data represented as x□±SEM.

After adding adolescence COC intake as a covariate in the Two Way ANCOVAs the main group effects remained in ACC (F_(1,6)_=9.006, *p*=.024, η_p_^2^=.600), OFC (F_(1,6)_=10.699, *p*=.017, η_p_^2^=.641), SGC (F_(1,6)_=7.946, *p*=.030, η_p_^2^=.570), dlPFC (F_(1,6)_=16.104, *p*=.007, η_p_^2^=.729), and vmPFC (F_(1,6)_=16.371, *p*=.007, η_p_^2^=.732) but disappeared in NAcc although there was a trend towards significance (F_(1,6)_=4.782, *p*=.071, η_p_^2^=.444). In all these ROIs, MALT animals showed significantly lower 5HT1A Receptor BP than Controls. Main time effects remained in OFC (F_(1,6)_=7.146, *p*=.037, η_p_^2^=.544), SGC (F_(1,6)_=12.595, *p*=.012, η_p_^2^=.677), and dlPFC (F_(1,6)_=13.969, *p*=.010, η_p_^2^=.700) but disappeared in Hippocampus, mPFC, vlPFC, and vmPFC. However, vlPFC (F_(1,6)_=5.602, *p*=.056, η_p_^2^=.483) and vmPFC (F_(1,6)_=5.829, *p*=.052, η_p_^2^=.493) had a trend towards significance. In all these ROIs except OFC, animals showed higher 5HT_1A_ receptor BP post-COC SA than at baseline; OFC showed lower 5HT1A receptor BP post-COC SA than at baseline. Significant group x time interaction effects remained in OFC (F_(1,6)_=34.951, *p*=.001, η_p_^2^=.853), SGC (F_(1,6)_=23.887, *p*=.003, η_p_^2^=.799), dlPFC (F_(1,6)_=14.800, *p*=.008, η_p_^2^=.712), and vlPFC (F_(1,6)_=7.632, *p*=.033, η_p_^2^=.560).

#### 5HT2A Receptor *BP*

RM ANOVA revealed significant main effects of group in dlPFC (F_(1,7)_=7.037, *p*=.033, η_p_^2^=.501) and vmPFC (F_(1,7)_=9.433, *p*=.018, η_p_^2^=.574), with MALT animals showing lower 5HT_2A_ receptor BP than Controls (**Figure 3**). Additionally, significant main time effects were found in ACC (F_(1,7)_=20.651, *p*=.003, η_p_^2^=.747), NAcc (F_(1,7)_=8.782, *p*=.021, η_p_^2^=.556), OFC (F_(1,7)_=33.910, *p*=.001, η_p_^2^=.829), SGC (F_(1,7)_=16.414, *p*=.005, η_p_^2^=.701), dlPFC (F_(1,7)_=21.807, *p*=.002, η_p_^2^=.757), mPFC (F_(1,7)_=10.881, *p*=.013, η_p_^2^=.609), vlPFC (F_(1,7)_=8.939, *p*=.020, η_p_^2^=.561), and vmPFC (F_(1,7)_=14.356, *p*=.007, η_p_^2^=.672). In all these ROIs, animals showed higher 5HT_2A_ receptor BP post-COC SA than at baseline (**Figure 3**). No group x time interaction effects were detected.

**Figure 3.**
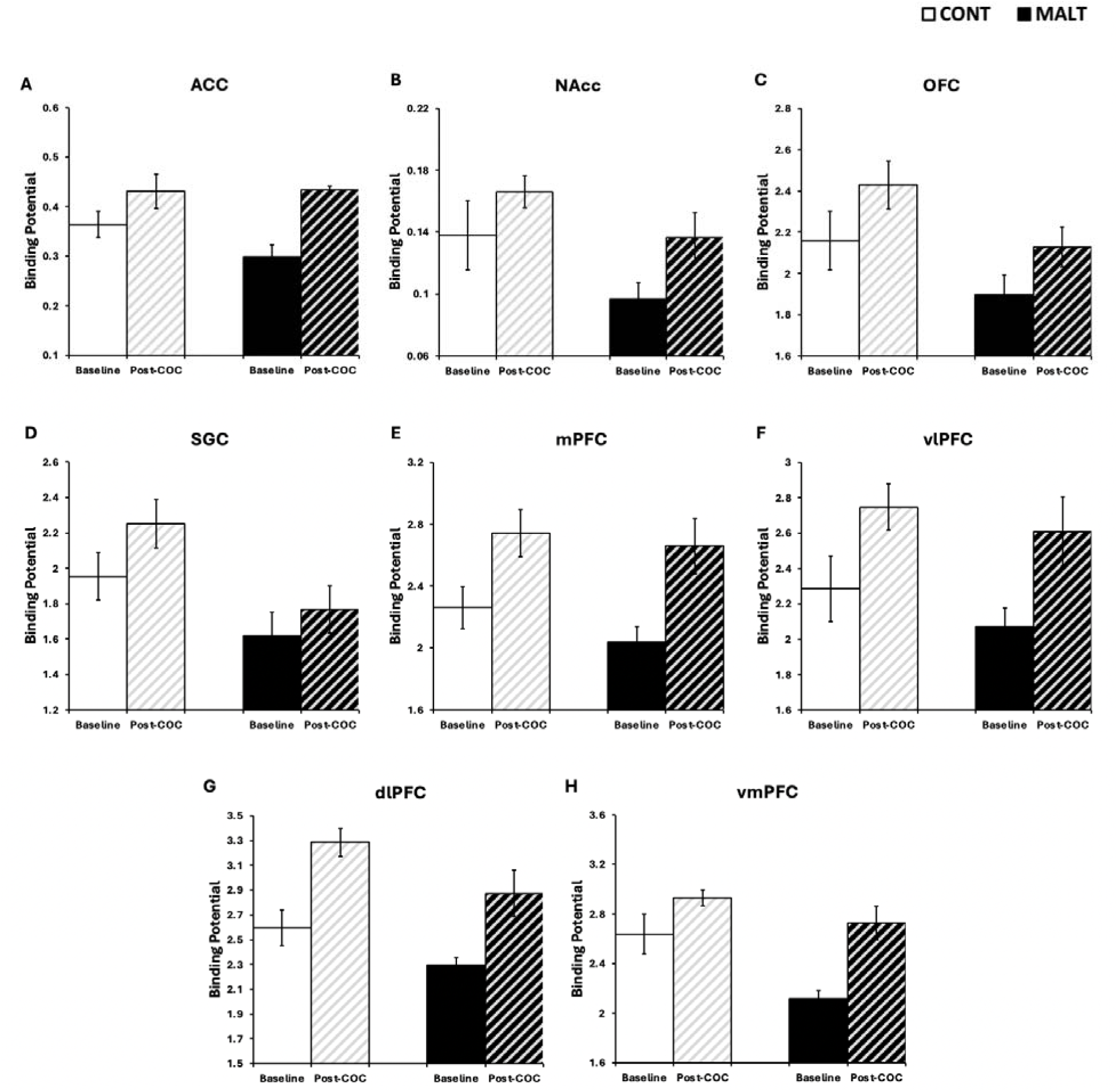
Changes in 5HT_2A_ receptor Binding Potential (BP) from Baseline (pre-cocaine[COC] self-administration [SA]) to post-COC SA in corticolimbic regions studied using [F18]-MDL100. Significant main group effects were detected in dlPFC and vmPFC, with lower 5HT_2A_ receptor BP in MALT than Control animals. Main time effects were also found in NAcc, ACC, OFC, SGC, dlPFC, mPFC, vmPFC and vlPFC, showing overall main receptor upregulation following COC SA, compared to baseline. No group x time interaction effects were found. See statistical details in text. NAcc: Nucleus Accumbens, ACC: anterior cingulate cortex, mPFC: medial prefrontal cortex; OFC: orbitofrontal Cortex, SGC: subgenual cingulate cortex, dlPFC: dorsolateral PFC, vlPFC: ventrolateral PFC, vmPFC: ventromedial PFC. Data represented as x□±SEM.

After adding adolescence COC intake as a covariate in the Two Way ANCOVAs the main group effects remained in dlPFC (F_(1,6)_=6.143, *p*=.048, η_p_^2^=.506), and vmPFC (F_(1,6)_=13.834, *p*=.010, η_p_^2^=.697) with MALT animals showing lower 5HT_2A_ receptor BP than Controls. However, the main time effect disappeared in the ACC, NAcc, OFC, SGC, dlPFC, mPFC, vlPFC, and vmPFC, although trend towards significance were still detected in the OFC (F_(1,6)_=4.190, *p*=.087, η_p_^2^=.411) and dlPFC (F_(1,6)_=4.631, *p*=.075, η_p_^2^=.436). Adolescence COC intake had a significant effect as covariate in NAcc (F_(1,6)_=9.410, *p*=0.022, η_p_^2^=.611) and vmPFC (F_(1,6)_=6.982, *p*=.038, η_p_^2^=.538).

#### D2/D 3 Receptor BP

RM ANOVA revealed significant time effects in Hippocampus (F_(1,7)_= 16.114, *p*=.005, η_p_^2^=.697), dlPFC (F_(1,7)_=30.392, *p*=.001, η_p_^2^=.813) and vlPFC (F_(1,7)_=8.370, *p*=.023, η_p_^2^=.545). In all these ROIs, higher D_2_/D_3_ receptor BP were found post-COC SA than at baseline (**Figure 4**).

**Figure 4.**
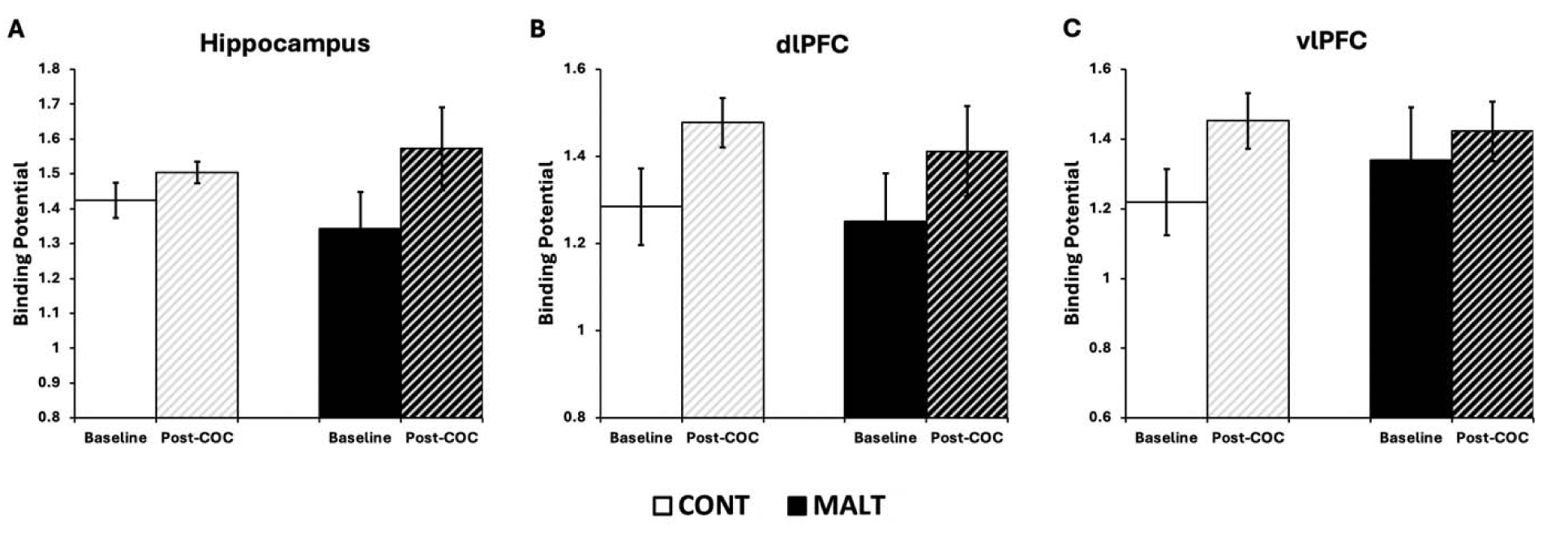
Changes in D_2_/D_3_ receptor binding potential (BP) from Baseline (pre-cocaine[COC] self-administration [SA]) to post-COC SA in corticolimbic regions studied using [F18]-fallypride. Main time effects were found in Hippocampus, dlPFC and vlPFC showing overall upregulation of D_2_/D_3_ receptor BP following COC SA, compared to baseline. No other main group, or group x time interaction effects were found. See statistical details in text. dlPFC: dorsolateral PFC, vlPFC: ventrolateral PFC. Data represented as x□±SEM.

After adding adolescence COC intake as a covariate in the Two Way ANCOVAs main time effects remained in Hippocampus (F_(1,6)_=11.196, *p*=.015, η_p_^2^=.651), but no longer in the dlPFC and vlPFC, although we detected a trend towards significance in the dlPFC (F(1,6) =5.014, *p*=0.066, η_p_^2^=.455). Animals showed higher D_2_/D_3_ receptor BP post-COC SA than at baseline. Adolescence COC intake had a significant effect as covariate in NAcc F(1,6) =14.421, *p*=.009, η_p_^2^=.706).

### Associations Between Adult Brain PET ROI BP and COC SA data

#### Identifying Networks of “Clustered” ROIs by PET Ligands

The Spearman Rank’s correlation significant clustering results are shown in correlation matrices for the different PET ligands in **Suppl. Tables 1-3**. Clusters varied by ligand and we generated averages of PET BP in ROIs included in the following clusters: (1) [18F]MPPF (**Suppl. Table 1**): Cluster 1 (dlPFC, mPFC, vmPFC) and Cluster 2 (ACC, OFC, SGC, mPFC), in addition to ROIs that did not cluster with others (Amygdala, Hippocampus, NAcc, Caudate, Putamen, vlPFC); (2) [18F]MDL100 (**Suppl. Table 2**): Cluster 1 (ACC, dlPFC, mPFC, vmPFC, and vlPFC), Cluster 2 (ACC, OFC), Cluster 3 (Amygdala, OFC, Caudate, Putamen), Cluster 4 (Amygdala, NAcc, OFC, SGC), and Cluster 5 (Hippocampus, ACC, Caudate, dlPFC, mPFC, vlPFC, vmPFC); and (3) [18F]fallypride (**Suppl. Table 3**): Cluster 1 (ACC, OFC, SGC, mPFC, dlPFC, vlPFC, vmPFC) and ROIs that did not cluster with others (Amygdala, Hippocampus, NAcc, Caudate, Putamen). These clusters were then entered into Spearman Rank correlation analyses of associations between PET and Adult COC SA data published recently (Allen et al., 2025). Adult COC SA data included in the models: (1) peak COC dose RR in the FR20 schedule, and (2) PR peak COC break point [note that no significant group or group x sex effects were detected on either of these measures (Allen et al., 2025)].

#### Spearman’s Rank Correlations Between Receptor BPs (ROI Clusters) and Adult COC SA Data

As shown in **Figure 5**, baseline Amygdala D2/D3 receptor BP was negatively correlated with FR RR (rho=-.632, *p*=.021), while 5HT_1A_ receptor BP in the same ROI was negatively correlated with PR peak Breakpoint (rho=.611, *p*=.046). No other significant associations were detected.

**Figure 5.**
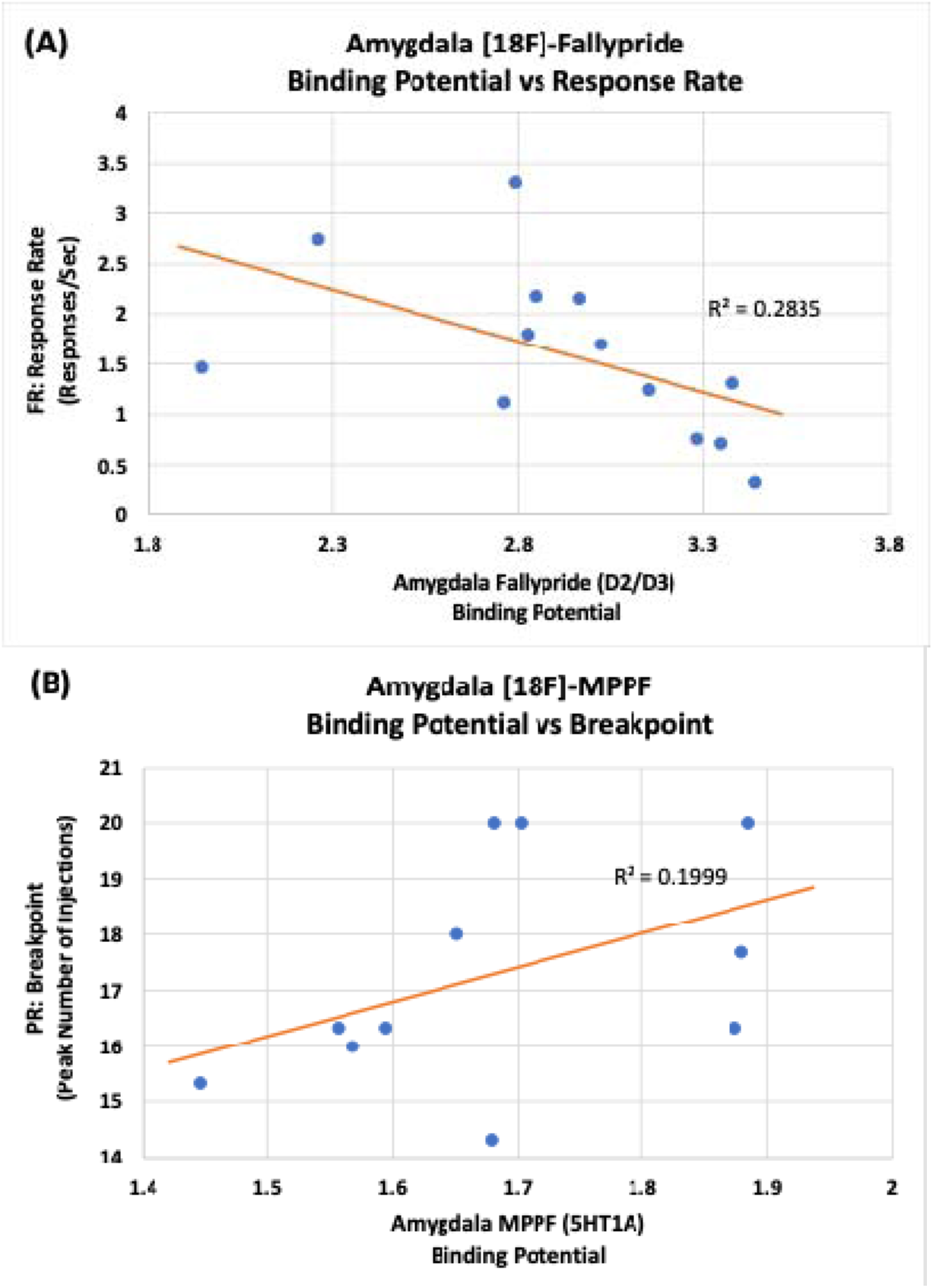
Regression Plots Based on Significant Spearman’s Rank Correlations Between Baseline PET (clusters) and COC SA data. (A) Amygdala baseline [18F]fallypride Binding Potential (BP) was negatively correlated with COC SA FR peak dose Response Rate (rho=-.632, *p*=.021). (B) Amygdala baseline [18F]MPPF BP was positively correlated with COC SA PR peak dose Breakpoint (rho=.611, *p*=.046).

## Discussion

This study examined the long-term effects of ELA on adult 5HT and DA function in corticolimbic regions important for reward and stress/emotional regulation, using a translational infant MALT macaque model and COC SA. The study focused on regional 5HT_1A_, 5HT_2A_, and D_2_/D_3_ receptor availability (BP) differences between MALT and Control animals using PET, both at baseline (pre-COC SA) and in response to chronic COC exposure. Additionally, we examined whether levels of these 5HT and DA receptors predicted COC SA outcomes, in particular COC reinforcing effects and potency, using FR peak RR and PR peak breakpoint. Through our results, long-term effects of infant MALT on 5HT, but not DA, receptors in corticolimbic circuits were apparent. Specifically, MALT animals showed lower 5HT_1A_ BP in the ACC, mPFC, and hippocampus compared to Controls. A MALT by Sex interaction effect was detected in 5HT_2A_ BP in the OFC, with lower levels in MALT than Control males, but not in females. In addition, upregulation of 5HT_1A_ and 5HT_2A_ receptors following chronic COC SA was detected in most PFC subregions, hippocampus, and NAcc, particularly in Control animals. Interestingly, we did not detect the expected downregulation in D_2_/D_3_ receptors in NAcc, caudate or putamen previously reported in the literature; in fact, we found an upregulation of D_2_/D_3_ receptors in dlPFC, vlPFC and hippocampus. We also found associations between PET baseline receptor BP data and COC SA measures; specifically, amygdala baseline levels of 5HT_1A_ receptors were positively correlated with PR peak breakpoint, while a negative correlation was found between baseline D2/D3 receptors and FR RR. Altogether these findings suggest broad long-term effects of ELA on adult brain 5HT, but not DA, receptors in corticolimbic regions involved in emotional and reward processes. We also found upregulation of 5HT_1A_ and 5HT_2A_ receptors in several corticolimbic regions following chronic COC exposure, suggesting an important role of 5HT receptors in early COC stages, as well as a role of the amygdala D2/D3 and 5HT_1A_ receptor availability on reward motivated behaviors. The dynamic range of changes of these 5HT_1A_ and 5HT_2A_ receptors following COC exposure was blunted in animals with ELA. It would be important to test whether the lack of dynamic range in these receptors could serve as biomarkers for CUD and can inform future pharmacotherapies that target 5HT receptors.

Lower 5HT_1A_ BP were detected in MALT subjects compared to Controls in PFC subregions responsible for stress/emotional regulation and connections with reward networks -ACC, mPFC- (Wakeford et al., 2024; Xu et al., 2019; Yalcin et al., 2014), as well as the hippocampus, an area involved with memory formation and consolidation (Zhu et al., 2021). Furthermore, MALT males exhibited lower 5HT_2A_ BP in the OFC compared to Controls, an area critical for goal-directed behavior and decision making, while no differences were detected for females (Hiser & Koenigs, 2018). These findings suggest long-term effects of ELA on adult 5HT, but not DA, receptors, with broad reductions in corticolimbic regions involved in emotional and reward processes, as well as in regions critical for executive function, cognitive control and memory (Hathaway & Newton, 2023; Wakeford et al., 2024; Zhu et al., 2021). This is consistent with previous reports in macaques showing that other ELA experiences such as maternal deprivation are linked to long-term alterations in serotonergic systems, including lower 5HT_1A_ receptor availability in brain regions involved in emotional regulation, stress responses, and executive function (Spinelli et al., 2010).

Our findings suggest persistent alterations in brain 5HT_1A_ and 5HT_2A_ and D_2_/D_3_ receptors in adults compared to adolescence in the same cohort of animals, where we showed that infant MALT led to reduced 5HT_1A_ receptor BP in PFC areas (ACC, SGC, dlPFC, mPFC, vlPFC, vmPFC), amygdala, and hippocampus compared to Controls, along with lower 5HT_2A_ receptor BP in PFC (OFC, dlPFC, vlPFC, vmPFC), NAcc and putamen in adolescent MALT macaques compared to Controls (Wakeford et al., 2024). ACC, mPFC, and hippocampus continue to show lower 5HT_1A_ receptor BP in adult MALT than Controls while other ROIs seemed to have normalized with age (SGC, dlPFC, vlPFC, vmPFC, amygdala). Lower 5HT_2A_ receptor BP persisted in adult MALT compared to Controls only in the OFC, while other ROIs normalized with age (dlPFC, vlPFC, vmPFC, NAcc, putamen). Long-term, region-specific effects of ELA on adult brain 5HT receptors have been reported in other studies. For example, adult rats exposed to ELA (early deprivation, maternal deprivation) show reduced 5HT_1A_ receptor binding in ACC, hippocampus, and PFC (Leventopoulos et al., 2009; Rentesi et al., 2013). Chronic or repeated stress in adult rodents also result in decreased levels of 5HT_1A_ (and sometimes 5HT_2A_) receptor levels in corticolimbic regions, specifically in the hippocampus. (Berton et al., 1998; Dwivedi et al., 2005; López et al., 1998; Mckittrick et al., 1995; Watanabe et al., 1993). The lower 5HT_1A_ BP in MALT than Controls in the NAcc, suggests a potential impact of these receptors in an area involved in reward processing (Xu et al., 2019); the lower levels in PFC subregions (ACC, OFC, SGC, dlPFC, and vmPFC) could imply roles of these 5HT receptors on top-down control on stress, emotional responses, and reward (Girotti et al., 2018; Hiser & Koenigs, 2018; Joyce et al., 2020; Joyce et al., 2024; Yalcin et al., 2014). To our knowledge, our study is the first of its kind using a NHP model of ELA to study longitudinally its long-term effects on 5HT and DA receptor function from adolescence to adulthood, in males and females. The fact that some 5HT_1A_ and 5HT_2A_ receptor alterations persist but others normalize from adolescence to adulthood in animals with ELA suggest some of them are “transient”. This is consistent with a recent publication by our group reporting similar transient ELA effects on hypothalamic-pituitary-adrenal (HPA) axis hyperactivity of MALT animals in this cohort, which normalized by adolescence (McCormack et al., 2025). This suggests that some ELA-induced neurochemical and neuroendocrine alterations can be rescued with time if the adversity stops. It is well-established that disruptions to the serotonergic system, including alterations in 5HT receptor binding, are associated with a heightened risk for mood and anxiety disorders such as depression, social anxiety, and panic disorder (Arango et al., 2001; Hirvonen et al., 2008; Lanzenberger et al., 2007; Nash et al., 2008), as well as impaired decision-making that contribute to maladaptive behavioral patterns (Adams et al., 2017). The long-lasting disruptions in 5HT receptor availability in adult MALT macaques suggest persistent neurobiological vulnerabilities consistent with increased risk for psychiatric disorders.

We observed lower baseline D_2_/D_3_ BP in males compared to females in the vlPFC. Given the vlPC’s role as a mediator between motor and sensory areas while receiving emotional and motivational information from the OFC and amygdala (Barbas & Deolmos, 1990; Ilinsky et al., 1985), these findings align with prior literature suggesting that sex differences in DA signaling in frontal regions. For example, in humans males exhibit lower extrastriatal D_2_-like receptor BP than females in regions such as the ACC, mPFC and dlPFC (Kaasinen et al., 2001). Similarly, lower DA receptor expression has been reported in male NHPs in caudate nucleus and in the rat striatum (Hasbi et al., 2020). These sex differences in DA signaling have been linked to sex specific difference in DA-related behaviors in neuropsychiatric disorders, including higher risk for drug addiction, including to psychostimulants like COC. Men with moderate nicotine addiction have lower D_2_ receptor availability in the striatum compared to females (Brown et al., 2012). Male mice undergoing chronic COC administration had less D_2_ receptor expressing neurons in the mPFC and the VTA compared to females (Clare et al., 2021). Although studies of sex differences in neuropsychiatric disorders that involve the vlPFC are scarce, dopaminergic dysfunction in neighboring regions, such as the OFC and amygdala play a role in SUD (Barbas & Deolmos, 1990; Ilinsky et al., 1985). For example, conditioned COC craving in women with SUD result in less neural activation of both the OFC and the amygdala (Kilts et al., 2004) than in COC-dependent men. What does this all mean? Maybe lower D_2_/D_3_ receptors in vlPFC, OFC and amygdala play a more important role in vulnerability of males to substance-induced behaviors such as stimulant craving and compulsive drug taking.

A striking upregulation of 5HT_1A_ and 5HT_2A_ receptors following chronic COC SA was detected in NAcc, most PFC subregions and hippocampus. Controls also showed higher levels of these 5HT receptors in dlPFC, vmPFC, ACC, OFC, SGC, vmPFC and NAcc than MALT animals. The dynamic response of both 5HT receptors to chronic COC exposure was also blunted in MALT animals. It would be important to examine whether the magnitude of change in these receptors BP could serve as biomarkers of CUD outcomes and inform future pharmacotherapies targeting 5HT receptors. During COC SA, 5HT receptor expression increases in the NAcc after using a viral-mediated 5HT receptor gene transfer (Pentkowski et al., 2014), which might magnify the role of 5HT modulating DA-mediated behavioral sensitization, therefore increasing stimulant consumption, which has been a mechanism reported at least during chronic stress exposure (Miczek et al., 2008).

Although speculative, the upregulation in both the 5HT_1A_ and 5HT_2A_ BP post COC SA suggest a possible compensatory adaptation to chronic COC exposure; particularly the upregulation of the inhibitory 5HT1A receptors in PFC regions projecting to the NAcc (e.g. mPFC, ACC), which would reduce COC reinforcing effects. Explaining how lower levels of 5HT receptors due to ELA may play a role in risk for SUD, is more difficult, unless it was the 5HT_2A_ receptor (excitatory). Conversely, low 5HT_1A_ and 5HT_2A_ receptor BP across PFC subregions and limbic areas of MALT animals, together with the blunted upregulation we uncovered following chronic COC SA, may indicate impaired plasticity of these receptors as a result of ELA. Previous studies support this possibility. For example, genetic deletion of 5HT_1A_ receptors leads to enhanced anxiety behaviors without altering 5HT levels and increasing 5HT turnover—possibly due to heightened serotonergic neuron activity or compensatory changes due to the lack of the receptors (Ase et al., 2000; Ramboz et al., 1998). In fact, introducing SSRIs to 5HT_1A_ knockout mice increases 5HT levels in the frontal cortex to a greater extent compared to controls, suggesting the absence of an inhibitory feedback control over 5HT release (Bortolozzi et al., 2004; Guilloux et al., 2006; Knobelman et al., 2001). Furthermore, individuals expressing polymorphism C(-1019)G results in not only alterations in levels of 5HT_1A_ receptor expression, but also influences susceptibility to stress and is associated with SUD (Albert & Lemonde, 2004; Huang et al., 2004). This evidence supports the idea that serotonergic plasticity may be crucial as a form of compensatory mechanism, protecting against the effects of chronic drug and stress exposure. One region that stood out was the SGC, where, following COC SA, Control animals showed increased 5HT_1A_ receptor BP but no change in the MALT group. The SGC (area 25) contains very high 5HT_1A_ receptor densities (Palomero-Gallagher et al., 2009; Varnäs et al., 2004) and has strong projections to the dorsal raphe (Freedman et al., 2000), source of serotoninergic projections to most of the cerebral cortex as well as the NAcc and VTA (Howell & Cunningham, 2015). Given the inhibitory nature of this receptor, area 25 could play a vital role in the inhibition of 5HT neurotransmission and, therefore, an effect on DA signaling in other areas. Although speculative, the lack of 5HT_1A_ receptor plasticity in MALT animals in contrast to the upregulation in Controls following COC SA, may indicate a different physiological response specific to this group, making area 25 a critical target. An alternative explanation of our findings is that the lower 5HT_1A_ receptor levels in MALT animals are an adaptation to increase DA neurotransmission, and therefore hedonic tone in the animals, which would be consistent with reports of anhedonia in rodent ELA models (Bangasser & Cuarenta, 2021; Bolton et al., 2017; Bolton et al., 2018; Sanchez et al., 2022).

Upregulation in D_2_/D_3_ BP following chronic COC compared to baseline was detected in dlPFC, vlPFC and hippocampus, without MALT or interaction effects. This result was puzzling, as previous studies have reported D_2_/D_3_ receptor downegulation in NAcc, caudate and putamen in macaques chronically exposed to COC (Allen et al., 2023), interpreted as a compensatory response to the increased in DA in the synapse due to COC blocking DAT reuptake (Bolla et al., 1998). One possible explanation is the lower COC exposure in our study (100 mgCOC/kg Bodyweight) compared to higher total COC intake in other studies (e.g. 300 mgCOC/kg; (Allen et al., 2023). Altogether, these patterns of r5HT and DA receptor upregulation may reflect neuroadaptive interactive mechanisms (Muller et al., 2007) in the early onset phase of COC use to counteract maladaptive COC effects (Howell & Cunningham, 2015). To our knowledge, this is the first study to demonstrate neurochemical changes that occur after chronic COC use, providing insight into the early progression of cocaine-induced alterations and potential targets for treatment.

ELA interferes with normal development of specific brain regions such as the hippocampus and amygdala, both involved in the stress response, as well as the mesocorticolimbic DA pathway which is closely tied to sensitivity and reward processing (Guyer et al., 2006; Kirsch & Lippard, 2022; Teicher & Samson, 2016). ELA also affects sensitivity to the levels of drug-induced euphoria (Wand et al., 2007), with evidence suggesting that stronger DA responses to substance use could increase the reinforcing properties of drugs (Kirsch & Lippard, 2022). For example, individuals with a history of childhood adversity show greater DA response in ventral striatum than healthy controls to amphetamine SA (Oswald et al., 2014). Individuals with mood and anxiety disorders—which frequently co-occur with SUDs— often report that addictive substances, such as alcohol, enhances euphoric mood (Kirsch & Lippard, 2022; McDonald & Meyer, 2011; Norberg et al., 2010). Given a high comorbidity between and mood and anxiety disorders, a history of ELA may contribute to heightened sensitivity to addictive substances, increasing risk for compulsive drug use at lower amounts.

We also examined associations between baseline PET receptor BP data and COC SA measures in adults. From the brain region clusters, we first found a positive correlation between baseline 5HT_1A_ PET receptor BP and PR Breakpoint in the amygdala, which suggests increased motivation/effort to obtain COC with higher receptor availability. This is consistent with the inhibitory role of this receptor and evidence that 5HT agonists that reduce the firing rate of mesolimbic DA neurons, reducing drug seeking behaviors (Fletcher et al., 2008). In addition, a negative correlation was detected between baseline D_2_/D_3_ PET receptor BP in the amygdala and COC SA FR response rates, suggesting that DA signaling in this region may modulate COC reinforcing effects in a schedule-dependent manner via amygdala’s direct projections to the NAcc (Stuber et al., 2011) and its role in reward processing. Glutamatergic projections from the basolateral amygdala (BLA) to the NAcc modulate cue-triggered motivated behaviors (Fareri & Tottenham, 2016; Stuber et al., 2011). DA input to the amygdala acting on D_2_/D_3_ receptors could inhibit these glutamatergic interactions and reduce reward behaviors by decreasing the reinforcing effects of drugs (Diaz et al., 2011; Fareri & Tottenham, 2016).

Several limitations of the current study need to be considered when interpreting the findings. First, although the sample size was large for this type of macaque studies and sufficient for the analysis of group effects on baseline brain PET data, it is underpowered to detect some of the interaction effects in the RM ANOVA, particularly in the studies of D_2_/D_3_, 5HT_1A_ and 5HT_2A_ receptor changes following chronic COC exposure. Given previous reports in the literature highlighting sex differences in risk for SUDs, including in populations with ELA experience (Brown et al., 2012; Clare et al., 2021; Kilts et al., 2004), and our previous reports of sex differences when these animals were adolescents (Wakeford et al., 2024), future studies should be done with bigger sample sizes. Second, some of the findings suggesting long-lasting neurobiological effects of ELA (MALT) on DA and 5HT receptors in adulthood might have been influenced by adolescent COC exposure. Adolescence exposure to psychostimulants affects the development of mesocortical and mesostriatal DA pathways (Pantoja-Urbán et al., 2024; Vassilev et al., 2021) and could have confounded the ELA effects on DA and 5HT receptors, affecting behavior. Indeed, a current publication by our lab (Allen et al., 2025) showed that COC SA FR RR are higher in our adults exposed to higher total COC intake during adolescence. However, another study that examined the rate of recovery of D_2_ receptors in rhesus macaques who underwent at least a year of COC SA showed full recovery of D_2_ receptor availability within three months of abstinence (Nader et al., 2006). Despite this evidence of receptor recovery, it remains unclear whether there are other receptor alterations beyond DA receptors (e.g. for 5HT receptors) from prior cocaine use. Although we entered total adolescence COC intake as a covariate in the statistical models, future studies should address the long-term effects of ELA with adult COC naïve animals. Third, the correlations between baseline PET receptor BP and COC SA measures, required data reduction for the predictors (baseline PET receptor BP), particularly given our small sample size. For this, a correlation-based clustering method was used to identify clusters of associations between ROIs BPs for each PET ligand, using Spearman rank’s correlations followed by visualization of the significant clustering results in correlation matrices. Although acceptable, this might not be the best approach, as there are other data-driven classification methods such as Cluster Analysis, Hierarchical Clustering, Centroid-based Clustering that would have been more appropriate. Creating clusters of brain regions was necessary due to the limited number of subjects available for each outcome (n=13 for the response rate and n=11 for the peak breakpoint). Future studies with larger sample sizes would allow for more detailed analyses, potentially being able to individually evaluate separate brain regions in relation to the selected outcomes.

Despite the limitations, our findings are a major contribution to the literature on the long-term effects of ELA (MALT) on adult brain 5HT and DA receptor systems involved in COC reinforcement. Currently there are no FDA-approved treatments specific for CUD and current studies have had a primary focus on manipulating DA receptors (Schwartz et al., 2022). Our findings suggest long-term effects of ELA (at least due to MALT) on adult brain 5HT, not DA, receptors in corticolimbic regions involved in emotional and reward processes, as well as following chronic COC exposure. This finding supports the well-established role of the 5HT system in modulating DA function and its plasticity under chronic COC use, as 5HT receptors are located widely across key reward and stress/emotional regulatory neurocircuits, including mesolimbic and mesocortical dopaminergic pathways originating in VTA (Doherty & Pickel, 2000), NAcc (Mengod et al., 1990), and projecting to NAcc and PFC subregions (Celada et al., 2013; Martín-Ruiz et al., 2001; Mengod et al., 1990; Pazos & Palacios, 1985).

The 5HT_1A_ receptors in the PFC are located on glutamatergic neurons projecting to the VTA and as a result, modulates DA release in reward related areas such as the NAcc (Schenk & Highgate, 2021). Although speculative, an increase in 5HT_1A_ receptor BP for both MALT and control subjects following COC SA may reduce excitatory signals from the PFC to the NAcc and VTA, reducing the reinforcing effects of COC via weaker DA response in the NAcc. However, since MALT effects on baseline levels of 5HT were detected mainly in the PFC, it may that the overall levels of receptor BP remain lower compared to controls due to lasting neurobiological effects of ELS. Studies show that anhedonia, defined as the reduced ability to feel pleasure (Craske et al., 2016), is a potential behavioral long-lasting outcome due to ELA such as childhood trauma (Fan et al., 2021; Germine et al., 2015). Thus, the biological systems of MALT animals may need heightened reward-related activation to gain similar feelings of pleasure and reward—possibly contributing to the reduced baseline 5HT_1A_ receptor BP levels, greater DA release in the NAcc, and potential increased susceptibly to addiction. This suggests that ELA specific neuroadaptations may prove to be maladaptive in the face of chronic drug use, altering both serotonergic and dopaminergic signaling by increasing the maltreated subjects’ vulnerability to substance use and decreasing their sensitivity to reward.

The associations found between baseline amygdala PET receptors BP and COC SA suggest that both 5HT and DA systems in the amygdala play a very important role modulating motivation for COC reinforcement. A recent study shows that the amygdala has an important role in reinforcement learning through its excitatory projections from the amygdala to NAcc (Costa et al., 2024). Specifically, these projections encode the value of exploring more informative (value based) options to induce directed exploration (Costa et al., 2024). This proves to be a strategy to tradeoff between addressing uncertainty but maximizing potential reward (Costa et al., 2024).

Given our findings, future studies should test implementations of pharmacological interventions targeting 5HT receptors in early COC use stages. In addition, investigating neurochemical changes during a prolonged period of abstinence following chronic COC would provide additional insight into the role of 5HT or DA receptors plasticity on outcomes. It would be important to test whether these dynamic changes serve as biomarkers for risk for CUD and can inform different pharmacological interventions. These 5HT receptors, more so than D_2_/D_3_ receptors, present promising targets through their ability to modulate DA signaling and circuit changes in early COC use stages.

## Author Contributions

Conceptualization: M.M. Sanchez, M.A. Nader, J.A. Nye

Methodology: J.A. Nye, M.A. Nader, E.R. Siebert, J. Khan, Ronald J. Voll, Lahu N. Chavan, Mark M. Goodman, J.H. Acevedo-Polo, M.M. Sanchez

Analyses: J.H. Acevedo-Polo, J.A. Nye, M.I. Rough, M.A. Nader, M.M. Sanchez

Data Curation and Visualization: J.H. Acevedo-Polo, J.A. Nye, M.M. Sanchez

Writing-Original Draft Preparation: J.H. Acevedo-Polo, J.A. Nye, M.M. Sanchez

Writing-Review and Editing: J.H. Acevedo-Polo, E.R. Siebert, J. Khan, M.I. Rough, R.J. Voll, L.N. Chavan, M.M. Goodman, J.A. Nye, M.A. Nader, M.M. Sanchez

Supervision: M.M. Sanchez

Funding Acquisition: M.M. Sanchez, M. Nader

All authors have read and agreed to the published version of the manuscript.

## Ethics Statement

The animal study protocol was approved by the Emory Institutional Animal Care and Use Committee (IACUC), Emory University, Atlanta, GA, USA, under the following protocols: All procedures were performed in accordance with the Animal Welfare Act and the U.S. Department of Health and Human Services “Guide for the Care and Use of Laboratory Animals” and the guidelines of the Declaration of Helsinki. All studies approved by the Emory Institutional Animal Care and Use Committee (IACUC), Emory University, Atlanta, GA, USA, under the following protocols: 296-2008Y, approved on 05/01/2009; YER-2001377-012015, approved on 01/20/2012; YER-2002956-112517 GA, approved on 11/25/2014; YER-4000084-ENTRPR-A, approved on 11/17/2017; PROTO202000091, approved on 11/06/2020 and renewed on 10/19/2023.

## Acknowledgments

The authors want to thank Brittany Howell, Anne Glenn, Christine Marsteller and Dora Guzman, and the staff at the Imaging Core and Emory National Primate Research Center (ENPRC) for their contributions, excellent technical support and animal care provided during these studies. In addition, we thank Dr. Melinda Higgins for Biostatistical guidance and support.

## Data Availability Statement

Dataset available on request from the authors. Data collection for the animals in this manuscript is ongoing for later endpoints.

## References

Adams, W. K., Barkus, C., Ferland, J. M. N., Sharp, T., & Winstanley, C. A. (2017). Pharmacological evidence that 5-HT receptor blockade selectively improves decision making when rewards are paired with audiovisual cues in a rat gambling task. Psychopharmacology, 234(20), 3091–3104. 10.1007/s00213-017-4696-4

Akimova, E., Lanzenberger, R., & Kasper, S. (2009). The serotonin-1A receptor in anxiety disorders. Biol Psychiatry, 66(7), 627–635. 10.1016/j.biopsych.2009.03.012

Albert, P. R., & Lemonde, S. (2004). 5-HT1A receptors, gene repression, and depression: Guilt by association. Neuroscientist, 10(6), 575–593. 10.1177/1073858404267382

Alexander, L., Wood, C. M., Gaskin, P. L. R., Sawiak, S. J., Fryer, T. D., Hong, Y. T., McIver, L., Clarke, H. F., & Roberts, A. C. (2020). Over-activation of primate subgenual cingulate cortex enhances the cardiovascular, behavioral and neural responses to threat. Nature Communications, 11(1). 10.1038/s41467-020-19167-0

Allen, M. I., Duke, A. N., Nader, S. H., Adler-Neal, A., Sai, K. K. S., Reboussin, B. A., Gage, H. D., Voll, R. J., Mintz, A., Goodman, M. M., & Nader, M. A. (2023). PET imaging of dopamine transporters and D2/D3 receptors in female monkeys: effects of chronic cocaine self-administration. Neuropsychopharmacology, 48(10), 1436–1445. 10.1038/s41386-023-01622-3

Allen, M. I., Siebert, E. R., Wakeford, A. G. P., Jenkins, K., Khan, J., Howell, L. L., Sanchez, M. M., & Nader, M. A. (2025). Cocaine self-administration in adult female and male rhesus monkeys: longitudinal comparison with adolescent behavior and role of early life stress. Neuropsychopharmacology, 50(13), 2006–2014. 10.1038/s41386-025-02161-9

Altmann, S. A. (1962). A field study of the sociobiology of rhesus monkeys, Macaca mulatta. Ann N Y Acad Sci, 102, 338–435. 10.1111/j.1749-6632.1962.tb13650.x

Arango, V., Underwood, M. D., Boldrini, M., Tamir, H., Kassir, S. A., Hsiung, S. C., Chen, J. J. X., & Mann, J. J. (2001). Serotonin 1A receptors, serotonin transporter binding and serotonin transporter mRNA expression in the brainstem of depressed suicide victims. Neuropsychopharmacology, 25(6), 892–903. 10.1016/S0893-133X(01)00310-4

Ase, A. R., Reader, T. A., Hen, R., Riad, M., & Descarries, L. (2000). Altered Serotonin and Dopamine Metabolism in the CNS of Serotonin 5-HT1A or 5-HT1B Receptor Knockout Mice. Journal of Neurochemistry, 75(6), 2415–2426. 10.1046/j.1471-4159.2000.0752415.x

Ayano, G. (2016). Dopamine: Receptors, Functions, Synthesis, Pathways, Locations and Mental Disorders: Review of Literatures. Journal of Mental Disorders and Treatment, 2. 10.4172/2471-271X.1000120

Balouek, J. A., Mclain, C. A., Minerva, A. R., Rashford, R. L., Bennett, S. N., Rogers, F. D., & Peña, C. J. (2023). Reactivation of Early-Life Stress-Sensitive Neuronal Ensembles Contributes to Lifelong Stress Hypersensitivity. Journal of Neuroscience, 43(34), 5996–6009. 10.1523/Jneurosci.0016-23.2023

Bangasser, D. A., & Cuarenta, A. (2021). Sex differences in anxiety and depression: circuits and mechanisms. Nature Reviews Neuroscience, 22(11), 674–684. 10.1038/s41583-021-00513-0

Barbas, H., & Deolmos, J. (1990). Projections from the Amygdala to Basoventral and Mediodorsal Prefrontal Regions in the Rhesus-Monkey. Journal of Comparative Neurology, 300(4), 549–571. 10.1002/cne.903000409

Barfield, E. T., Sequeira, M. K., Parsons, R. G., & Gourley, S. L. (2020). Morphological Responses of Excitatory Prelimbic and Orbitofrontal Cortical Neurons to Excess Corticosterone in Adolescence and Acute Stress in Adulthood. Frontiers in Neuroanatomy, 14. 10.3389/fnana.2020.00045

Berton, O., Aguerre, S., Sarrieau, A., Mormede, P., & Chaouloff, F. (1998). Differential effects of social stress on central serotonergic activity and emotional reactivity in Lewis and spontaneously hypertensive rats. Neuroscience, 82(1), 147–159. 10.1016/s0306-4522(97)00282-0

Bolla, K. I., Cadet, J. L., & London, E. D. (1998). The neuropsychiatry of chronic cocaine abuse. Journal of Neuropsychiatry and Clinical Neurosciences, 10(3), 280–289. 10.1176/jnp.10.3.280

Bolton, J. L., Molet, J., Ivy, A., & Baram, T. Z. (2017). New insights into early-life stress and behavioral outcomes. Current Opinion in Behavioral Sciences, 14, 133–139. 10.1016/j.cobeha.2016.12.012

Bolton, J. L., Molet, J., Regev, L., Chen, Y. C., Rismanchi, N., Haddad, E., Yang, D. Z., Obenaus, A., & Baram, T. Z. (2018). Anhedonia Following Early-Life Adversity Involves Aberrant Interaction of Reward and Anxiety Circuits and Is Reversed by Partial Silencing of Amygdala Corticotropin-Releasing Hormone Gene. Biological Psychiatry, 83(2), 137–147. 10.1016/j.biopsych.2017.08.023

Bortolozzi, A., Amargós-Bosch, M., Toth, M., Artigas, F., & Adell, A. (2004). In vivo efflux of serotonin in the dorsal raphe nucleus of 5-HT receptor knockout mice. Journal of Neurochemistry, 88(6), 1373–1379. 10.1046/j.1471-4159.2003.02267.x

Bronk, G., Lardenoije, R., Koolman, L., Klengel, C., Dan, S., Howell, B. R., Morin, E. L., Meyer, J. S., Wilson, M. E., Ethun, K. F., Alvarado, M. C., Raper, J., Bravo-Rivera, H., Kenwood, M. M., Roseboom, P. H., Quirk, G. J., Kalin, N. H., Binder, E. B., Sanchez, M. M., & Klengel, T. (2024). A novel epigenetic clock for rhesus macaques unveils an association between early life adversity and epigenetic age acceleration. bioRxiv. 10.1101/2024.10.08.617208

Brown, A. K., Mandelkern, M. A., Farahi, J., Robertson, C., Ghahremani, D. G., Sumerel, B., Moallem, N., & London, E. D. (2012). Sex differences in striatal dopamine D_2_/D_3_ receptor availability in smokers and non-smokers. International Journal of Neuropsychopharmacology, 15(7), 989–994. 10.1017/S1461145711001957

Camara, E., Rodriguez-Fornells, A., Ye, Z., & Münte, T. F. (2009). Reward networks in the brain as captured by connectivity measures. Frontiers in Neuroscience, 3(3), 350–362. 10.3389/neuro.01.034.2009

Carr, C. P., Martins, C. M. S., Stingel, A. M., Lemgruber, V. B., & Juruena, M. F. (2013). The Role of Early Life Stress in Adult Psychiatric Disorders. Journal of Nervous and Mental Disease, 201(12), 1007–1020. 10.1097/Nmd.0000000000000049

Celada, P., Puig, M. V., & Artigas, F. (2013). Serotonin modulation of cortical neurons and networks. Front Integr Neurosci, 7, 25. 10.3389/fnint.2013.00025

Chavan, L. N., Voll, R., Sanchez, M. M., Nye, J. A., & Goodman, M. M. (2023). Concise and Scalable Radiosynthesis of (+)-[^18^F]MDL100907 as a Serotonin 5-HT_2A_ Receptor Antagonist for PET. Acs Chemical Neuroscience, 14(19), 3694–3703. 10.1021/acschemneuro.3c00382

Christian, B. T., Vandehey, N. T., Fox, A. S., Murali, D., Oakes, T. R., Converse, A. K., Nickles, R. J., Shelton, S. E., Davidson, R. J., & Kalin, N. H. (2009). The distribution of D2/D3 receptor binding in the adolescent rhesus monkey using small animal PET imaging. Neuroimage, 44(4), 1334–1344. 10.1016/j.neuroimage.2008.10.020

Chu, D. A., Williams, L. M., Harris, A. W. F., Bryant, R. A., & Gatt, J. M. (2013). Early life trauma predicts self-reported levels of depressive and anxiety symptoms in nonclinical community adults: Relative contributions of early life stressor types and adult trauma exposure. Journal of Psychiatric Research, 47(1), 23–32. 10.1016/j.jpsychires.2012.08.006

Cicchetti, D., & Toth, S. L. (2005). Child maltreatment. Annual Review of Clinical Psychology, 1, 409–438. 10.1146/annurev.clinpsy.1.102803.144029

Clare, K., Pan, C., Kim, G., Park, K., Zhao, J., Volkow, N. D., Lin, Z. C., & Du, C. W. (2021). Cocaine Reduces the Neuronal Population While Upregulating Dopamine D2-Receptor-Expressing Neurons in Brain Reward Regions: Sex-Effects. Frontiers in Pharmacology, 12. 10.3389/fphar.2021.624127

Costa, V. D., Rothenhoefer, K. M., & Romac, M. D. (2024). Chemogenetic Inhibition of Amygdala Inputs to Striatum Modulates Reinforcement Learning. Poster presented at the 2024 Annual Meeting of the American College of Neuropsychopharmacology (ACNP), Tampa, FL.

Craske, M. G., Meuret, A. E., Ritz, T., Treanor, M., & Dour, H. J. (2016). Treatment for Anhedonia: A Neuroscience Driven Approach. Depression and Anxiety, 33(10), 927–938. 10.1002/da.22490

Diaz, M. R., Chappell, A. M., Christian, D. T., Anderson, N. J., & McCool, B. A. (2011). Dopamine D3-Like Receptors Modulate Anxiety-Like Behavior and Regulate GABAergic Transmission in the Rat Lateral/Basolateral Amygdala. Neuropsychopharmacology, 36(5), 1090–1103. 10.1038/npp.2010.246

Doherty, M. D., & Pickel, V. M. (2000). Ultrastructural localization of the serotonin 2A receptor in dopaminergic neurons in the ventral tegmental area. Brain Research, 864(2), 176–185. 10.1016/S0006-8993(00)02062-X

Douglas, K. R., Chan, G., Gelernter, J., Arias, A. J., Anton, R. F., Weiss, R. D., Brady, K., Poling, J., Farrer, L., & Kranzler, H. R. (2010). Adverse childhood events as risk factors for substance dependence: Partial mediation by mood and anxiety disorders. Addictive Behaviors, 35(1), 7–13. 10.1016/j.addbeh.2009.07.004

Drury, S. S., Howell, B. R., Jones, C., Esteves, K., Morin, E., Schlesinger, R., Meyer, J. S., Baker, K., & Sanchez, M. M. (2017). Shaping long-term primate development: Telomere length trajectory as an indicator of early maternal maltreatment and predictor of future physiologic regulation. Development and Psychopathology, 29(5), 1539–1551. 10.1017/S0954579417001225

Dunlop, B. W., & Nemeroff, C. B. (2007). The role of dopamine in the pathophysiology of depression. Arch Gen Psychiatry, 64(3), 327–337. 10.1001/archpsyc.64.3.327

Dwivedi, Y., Mondal, A. C., Payappagoudar, G. V., & Rizavi, H. S. (2005). Differential regulation of serotonin (5HT)2A receptor mRNA and protein levels after single and repeated stress in rat brain:: role in learned helplessness behavior. Neuropharmacology, 48(2), 204–214. 10.1016/j.neuropharm.2004.10.004

Embree, M., Michopoulos, V., Votaw, J. R., Voll, R. J., Mun, J., Stehouwer, J. S., Goodman, M. M., Wilson, M. E., & Sánchez, M. M. (2013). The Relation of Developmental Changes in Brain Serotonin Transporter (5htt) and 5ht1a Receptor Binding to Emotional Behavior in Female Rhesus Monkeys: Effects of Social Status and 5htt Genotype. Neuroscience, 228, 83–100. 10.1016/j.neuroscience.2012.10.016

Enoch, M. A. (2011). The role of early life stress as a predictor for alcohol and drug dependence. Psychopharmacology, 214(1), 17–31. 10.1007/s00213-010-1916-6

Everitt, B. J., & Robbins, T. W. (2005). Neural systems of reinforcement for drug addiction: from actions to habits to compulsion. Nature Neuroscience, 8(11), 1481–1489. 10.1038/nn1579

Fahlke, C., Lorenz, J. G., Long, J., Champoux, M., Suomi, S. J., & Higley, J. D. (2000). Rearing experiences and stress-induced plasma cortisol as early risk factors for excessive alcohol consumption in nonhuman primates. Alcoholism-Clinical and Experimental Research, 24(5), 644–650. 10.1097/00000374-200005000-00008

Fan, J., Liu, W. T., Xia, J., Li, S. H., Gao, F., Zhu, J., Han, Y., Zhou, H., Liao, H. Y., Yi, J. Y., Tan, C. L., & Zhu, X. Z. (2021). Childhood trauma is associated with elevated anhedonia and altered core reward circuitry in major depression patients and controls. Human Brain Mapping, 42(2), 286–297. 10.1002/hbm.25222

Fareri, D. S., & Tottenham, N. (2016). Effects of early life stress on amygdala and striatal development. Developmental Cognitive Neuroscience, 19, 233–247. 10.1016/j.dcn.2016.04.005

Fergusson, D. M., Boden, J. M., & Horwood, L. J. (2008). Exposure to childhood sexual and physical abuse and adjustment in early adulthood. Child Abuse & Neglect, 32(6), 607–619. 10.1016/j.chiabu.2006.12.018

Fletcher, P. J., Rizos, Z., Sinyard, J., Tampakeras, M., & Higgins, G. A. (2008). The 5-HT2C receptor agonist Ro60-0175 reduces cocaine self-administration and reinstatement induced by the stressor yohimbine, and contextual cues. Neuropsychopharmacology, 33(6), 1402–1412. 10.1038/sj.npp.1301509

Freedman, L. J., Insel, T. R., & Smith, Y. (2000). Subcortical projections of area 25 (subgenual cortex) of the macaque monkey. Journal of Comparative Neurology, 421(2), 172–188.

Fucich, E. A., Paredes, D., Saunders, M. O., & Morilak, D. A. (2018). Activity in the Ventral Medial Prefrontal Cortex Is Necessary for the Therapeutic Effects of Extinction in Rats. Journal of Neuroscience, 38(6), 1408–1417. 10.1523/Jneurosci.0635-17.2017

Gangopadhyay, P., Chawla, M., Dal Monte, O., & Chang, S. W. C. (2021). Prefrontal-amygdala circuits in social decision-making. Nature Neuroscience, 24(1), 5–18. 10.1038/s41593-020-00738-9

Germine, L., Dunn, E. C., McLaughlin, K. A., & Smoller, J. W. (2015). Childhood Adversity Is Associated with Adult Theory of Mind and Social Affiliation, but Not Face Processing. Plos One, 10(6). 10.1371/journal.pone.0129612

Girotti, M., Adler, S. M., Bulin, S. E., Fucich, E. A., Paredes, D., & Morilak, D. A. (2018). Prefrontal cortex executive processes affected by stress in health and disease. Progress in Neuro-Psychopharmacology & Biological Psychiatry, 85, 161–179. 10.1016/j.pnpbp.2017.07.004

Graves, F. C., & Wallen, K. (2006). Androgen-induced yawning in rhesus monkey females is reversed with a nonsteroidal anti-androgen. Horm Behav, 49(2), 233–236. 10.1016/j.yhbeh.2005.07.005

Greenfield, S. F., Kolodziej, M. E., Sugarman, D. E., Muenz, L. R., Vagge, L. M., He, D. Y., & Weiss, R. D. (2002). History of abuse and drinking outcomes following inpatient alcohol treatment: a prospective study. Drug and Alcohol Dependence, 67(3), 227–234. 10.1016/S0376-8716(02)00072-8

Guilloux, J. P., David, D. J., Guiard, B. P., Chenu, F., Repérant, C., Toth, M., Bourin, M., & Gardier, A. M. (2006). Blockade of 5-HT1A receptors by (+/−)-pindolol potentiates cortical 5-HT outflow, but not antidepressant-like activity of paroxetine: microdialysis and behavioral approaches in 5-HT1A receptor knockout mice. Neuropsychopharmacology, 31(10), 2162–2172. 10.1038/sj.npp.1301019

Guyer, A. E., Kaufman, J., Hodgdon, H. B., Masten, C. L., Jazbec, S., Pine, D. S., & Ernst, M. (2006). Behavioral alterations in reward system function: The role of childhood maltreatment and psychopathology. Journal of the American Academy of Child and Adolescent Psychiatry, 45(9), 1059–1067. 10.1097/01.chi.0000227882.50404.11

Hamani, C., Mayberg, H., Stone, S., Laxton, A., Haber, S., & Lozano, A. M. (2011). The Subcallosal Cingulate Gyrus in the Context of Major Depression. Biological Psychiatry, 69(4), 301–308. 10.1016/j.biopsych.2010.09.034

Hasbi, A., Nguyen, T., Rahal, H., Manduca, J. D., Miksys, S., Tyndale, R. F., Madras, B. K., Perreault, M. L., & George, S. R. (2020). Sex difference in dopamine D1-D2 receptor complex expression and signaling affects depression- and anxiety-like behaviors. Biology of Sex Differences, 11(1). 10.1186/s13293-020-00285-9

Hathaway, W. R., & Newton, B. W. (2023). Neuroanatomy, Prefrontal Cortex. In StatPearls. StatPearls Publishing.

Heffner, J. L., Blom, T. J., & Anthenelli, R. M. (2011). Gender Differences in Trauma History and Symptoms as Predictors of Relapse to Alcohol and Drug Use. American Journal on Addictions, 20(4), 307–311. 10.1111/j.1521-0391.2011.00141.x

Higley, J. D., Hasert, M. F., Suomi, S. J., & Linnoila, M. (1991). Nonhuman Primate Model of Alcohol-Abuse - Effects of Early Experience, Personality, and Stress on Alcohol-Consumption. Proceedings of the National Academy of Sciences of the United States of America, 88(16), 7261–7265. 10.1073/pnas.88.16.7261

Hirvonen, J., Karlsson, H., Kajander, J., Lepola, A., Markkula, J., Rasi-Hakala, H., Någren, K., Salminen, J. K., & Hietala, J. (2008). Decreased brain serotonin 5-HT1A receptor availability in medication-naive patients with major depressive disorder: an in-vivo imaging study using PET and [carbonyl-11C]WAY-100635. International Journal of Neuropsychopharmacology, 11(4), 465–476. 10.1017/S1461145707008140

Hiser, J., & Koenigs, M. (2018). The Multifaceted Role of the Ventromedial Prefrontal Cortex in Emotion, Decision Making, Social Cognition, and Psychopathology. Biological Psychiatry, 83(8), 638–647. 10.1016/j.biopsych.2017.10.030

Howell, B. R., Ahn, M., Shi, Y. D., Godfrey, J. R., Hu, X. P., Zhu, H. T., Styner, M., & Sanchez, M. M. (2019). Disentangling the effects of early caregiving experience and heritable factors on brain white matter development in rhesus monkeys. Neuroimage, 197, 625–642. 10.1016/j.neuroimage.2019.04.013

Howell, B. R., McCormack, K. M., Grand, A. P., Sawyer, N. T., Zhang, X., Maestripieri, D., Hu, X., & Sanchez, M. M. (2013). Brain white matter microstructure alterations in adolescent rhesus monkeys exposed to early life stress: associations with high cortisol during infancy. Biol Mood Anxiety Disord, 3(1), 21. 10.1186/2045-5380-3-21

Howell, B. R., McMurray, M. S., Guzman, D. B., Nair, G., Shi, Y., McCormack, K. M., Hu, X., Styner, M. A., & Sanchez, M. M. (2017). Maternal buffering beyond glucocorticoids: impact of early life stress on corticolimbic circuits that control infant responses to novelty. Soc Neurosci, 12(1), 50–64. 10.1080/17470919.2016.1200481

Howell, B. R., & Sanchez, M. M. (2011). Understanding behavioral effects of early life stress using the reactive scope and allostatic load models. Development and Psychopathology, 23(4), 1001–1016. 10.1017/S0954579411000460

Howell, L. L., & Cunningham, K. A. (2015). Serotonin 5-HT2 receptor interactions with dopamine function: implications for therapeutics in cocaine use disorder. Pharmacol Rev, 67(1), 176–197. 10.1124/pr.114.009514

Huang, Y. Y., Battistuzzi, C., Oquendo, M. A., Harkavy-Friedman, J., Greenhill, L., Zalsman, G., Brodsky, B., Arango, V., Brent, D. A., & Mann, J. J. (2004). Human 5-HT1A receptor C(-1019)G polymorphism and psychopathology. International Journal of Neuropsychopharmacology, 7(4), 441–451. 10.1017/S1461145704004663

Hyman, S. M., Paliwal, P., & Sinha, R. (2007). Childhood maltreatment, perceived stress, and stress-related coping in recently abstinent cocaine dependent adults. Psychology of Addictive Behaviors, 21(2), 233–238. 10.1037/0893-164x.21.2.233

Ilinsky, I. A., Jouandet, M. L., & Goldmanrakic, P. S. (1985). Organization of the Nigrothalamocortical System in the Rhesus-Monkey. Journal of Comparative Neurology, 236(3), 315–330. 10.1002/cne.902360304

Jaycox, L. H., Ebener, P., Damesek, L., & Becker, K. (2004). Trauma exposure and retention in adolescent substance abuse treatment. Journal of Traumatic Stress, 17(2), 113–121. 10.1023/B:Jots.0000022617.41299.39

Joyce, M. K. P., García-Cabezas, M. A., John, Y. J., & Barbas, H. (2020). Serial Prefrontal Pathways Are Positioned to Balance Cognition and Emotion in Primates. Journal of Neuroscience, 40(43), 8306–8328. 10.1523/Jneurosci.0860-20.2020

Joyce, M. K. P., Uchendu, S., & Arnsten, A. F. T. (2024). Stress and Inflammation Target Dorsolateral Prefrontal Cortex Function: Neural Mechanisms Underlying Weakened Cognitive Control. Biol Psychiatry. 10.1016/j.biopsych.2024.06.016

Kaasinen, V., Någren, K., Hietala, J., Farde, L., & Rinne, J. O. (2001). Sex differences in extrastriatal dopamine D_2_-like receptors in the human brain. American Journal of Psychiatry, 158(2), 308–311. 10.1176/appi.ajp.158.2.308

Kilts, C. D., Gross, R. E., Ely, T. D., & Drexler, K. P. G. (2004). The neural correlates of cue-induced craving in cocaine-dependent women. American Journal of Psychiatry, 161(2), 233–241. 10.1176/appi.ajp.161.2.233

Kirsch, D. E., & Lippard, E. T. C. (2022). Early life stress and substance use disorders: The critical role of adolescent substance use. Pharmacol Biochem Behav, 215, 173360. 10.1016/j.pbb.2022.173360

Klein, M. O., Battagello, D. S., Cardoso, A. R., Hauser, D. N., Bittencourt, J. C., & Correa, R. G. (2019). Dopamine: Functions, Signaling, and Association with Neurological Diseases. Cellular and Molecular Neurobiology, 39(1), 31–59. 10.1007/s10571-018-0632-3

Knobelman, D. A., Hen, R., Blendy, J. A., & Lucki, I. (2001). Regional patterns of compensation following genetic deletion of either 5-hydroxytryptamine(1A) or 5-hydroxytryptamine(1B) receptor in the mouse. J Pharmacol Exp Ther, 298(3), 1092–1100.

Koob, G. F., & Volkow, N. D. (2010). Neurocircuitry of Addiction. Neuropsychopharmacology, 35(1), 217–238. 10.1038/npp.2009.110

Kosten, T. A., Miserendino, M. J. D., & Kehoe, P. (2000). Enhanced acquisition of cocaine self-administration in adult rats with neonatal isolation stress experience. Brain Research, 875(1-2), 44–50. 10.1016/S0006-8993(00)02595-6

Kovacs-Balint, Z. A., Raper, J., Richardson, R., Gopakumar, A., Kettimuthu, K. P., Higgins, M., Feczko, E., Earl, E., Ethun, K. F., Li, L., Styner, M., Fair, D., Bachevalier, J., & Sanchez, M. M. (2023). The role of puberty on physical and brain development: A longitudinal study in male Rhesus Macaques. Developmental Cognitive Neuroscience, 60. 10.1016/j.dcn.2023.101237

Lammertsma, A. A., Bench, C. J., Hume, S. P., Osman, S., Gunn, K., Brooks, D. J., & Frackowiak, R. S. (1996). Comparison of methods for analysis of clinical [11C]raclopride studies. J Cereb Blood Flow Metab, 16(1), 42–52. 10.1097/00004647-199601000-00005

Lanzenberger, R. R., Mitterhauser, M., Spindelegger, C., Wadsak, W., Klein, N., Mien, L. K., Holik, A., Attarbaschi, T., Mossaheb, N., Sacher, J., Geiss-Granadia, T., Kletter, K., Kasper, S., & Tauscher, J. (2007). Reduced serotonin-1A receptor binding in social anxiety disorder. Biological Psychiatry, 61(9), 1081–1089. 10.1016/j.biopsych.2006.05.022

Lardenoije, R., Smulders, M. N. C. A., Morin, E. L., Howell, B. R., Guzman, D., Meyer, J. S., Ressler, K. J., Sanchez, M. M., & Klengel, T. (2025). A cross-generational methylomic signature of infant maltreatment in newborn rhesus macaques. Biological Psychiatry, In press, In press.

Le Bars, D., Lemaire, C., Ginovart, N., Plenevaux, A., Aerts, J., Brihaye, C., Hassoun, W., Leviel, V., Mekhsian, P., Weissmann, D., Pujol, J. F., Luxen, A., & Comar, D. (1998). High-yield radiosynthesis and preliminary in vivo evaluation of p-[18F]MPPF, a fluoro analog of WAY-100635. Nuclear Medicine and Biology, 25(4), 343–350. 10.1016/S0969-8051(97)00229-1

LeDoux, J. E. (1994). The amygdala: contributions to fear and stress. Seminars in Neuroscience, 6(4), 231–237. 10.1006/smns.1994.1030

Leventopoulos, M., Russig, H., Feldon, J., Pryce, C. R., & Opacka-Juffry, J. (2009). Early deprivation leads to long-term reductions in motivation for reward and 5-HT1A binding and both effects are reversed by fluoxetine. Neuropharmacology, 56(3), 692–701. 10.1016/j.neuropharm.2008.12.005

López, J. F., Chalmers, D. T., Little, K. Y., & Watson, S. J. (1998). Regulation of Serotonin1A, Glucocorticoid, and Mineralocorticoid Receptor in Rat and Human Hippocampus: Implications for the Neurobiology of Depression. Biological Psychiatry, 43(8), 547–573. 10.1016/S0006-3223(97)00484-8

Maes, F., Collignon, A., Vandermeulen, D., Marchal, G., & Suetens, P. (1997). Multimodality image registration by maximization of mutual information. Ieee Transactions on Medical Imaging, 16(2), 187–198. 10.1109/42.563664

Maestripieri, D. (1998). Parenting styles of abusive mothers in group-living rhesus macaques. Animal Behaviour, 55, 1–11. 10.1006/anbe.1997.0578

Maestripieri, D., & Carroll, K. A. (1998). Child abuse and neglect: Usefulness of the animal data. Psychological Bulletin, 123(3), 211–223. 10.1037/0033-2909.123.3.211

Maestripieri, D., Higley, J. D., Lindell, S. G., Newman, T. K., McCormack, K. M., & Sanchez, M. M. (2006). Early maternal rejection affects the development of monoaminergic systems and adult abusive parenting in rhesus macaques (Macaca mulatta). Behavioral Neuroscience, 120(5), 1017–1024. 10.1037/0735-7044.120.5.1017

Maestripieri, D., McCormack, K., Lindell, S. G., Higley, J. D., & Sanchez, M. M. (2006). Influence of parenting style on the offspring’s behaviour and CSF monoamine metabolite levels in crossfostered and noncrossfostered female rhesus macaques. Behavioural Brain Research, 175(1), 90–95. 10.1016/j.bbr.2006.08.002

Maestripieri, D., Megna, N. L., & Jovanovic, T. (2000). Adoption and maltreatment of foster infants by rhesus macaque abusive mothers. Developmental Science, 3(3), 287–293. 10.1111/1467-7687.00122

Mah, L., Szabuniewicz, C., & Fiocco, A. J. (2016). Can anxiety damage the brain? Curr Opin Psychiatry, 29(1), 56–63. 10.1097/yco.0000000000000223

Maier, S. F., & Watkins, L. R. (2010). Role of the medial prefrontal cortex in coping and resilience. Brain Research, 1355, 52–60. 10.1016/j.brainres.2010.08.039

Martín-Ruiz, R., Puig, M. V., Celada, P., Shapiro, D. A., Roth, B. L., Mengod, G., & Artigas, F. (2001). Control of serotonergic function in medial prefrontal cortex by serotonin-2A receptors through a glutamate-dependent mechanism. Journal of Neuroscience, 21(24), 9856–9866. 10.1523/Jneurosci.21-24-09856.2001

McCormack, K., Howell, B. R., Guzman, D., Villongco, C., Pears, K., Kim, H., Gunnar, M. R., & Sanchez, M. M. (2015). The Development of an Instrument to Measure Global Dimensions of Maternal Care in Rhesus Macaques (Macaca mulatta). American Journal of Primatology, 77(1), 20–33. 10.1002/ajp.22307

McCormack, K., Newman, T. K., Higley, J. D., Maestripieri, D., & Sanchez, M. M. (2009). Serotonin transporter gene variation, infant abuse, and responsiveness to stress in rhesus macaque mothers and infants. Hormones and Behavior, 55(4), 538–547. 10.1016/j.yhbeh.2009.01.009

McCormack, K., Sanchez, M. M., Bardi, M., & Maestripieri, D. (2006). Maternal care patterns and behavioral development of rhesus macaque abused infants in the first 6 months of life. Dev Psychobiol, 48(7), 537–550. 10.1002/dev.20157

McCormack, K. M., Bramlett, S., Morin, E. L., Siebert, E. R., Guzman, D., Howell, B. R., & Sanchez, M. M. (2025). Long-term effects of maternal care on hypothalamic-pituitary-adrenal (HPA) axis function of juvenile and adolescent macaques. Biology, In Press, In Press.

McCormack, K. M., Howell, B. R., Higgins, M., Bramlett, S., Guzman, D., Morin, E. L., Villongco, C., Liu, Y., Meyer, J., & Sanchez, M. M. (2022). The developmental consequences of early adverse care on infant macaques: A cross-fostering study. Psychoneuroendocrinology, 146, 105947. 10.1016/j.psyneuen.2022.105947

McDonald, J. L., & Meyer, T. D. (2011). Self-report reasons for alcohol use in bipolar disorders: why drink despite the potential risks? Clin Psychol Psychother, 18(5), 418–425. 10.1002/cpp.782

Mckittrick, C. R., Blanchard, D. C., Blanchard, R. J., Mcewen, B. S., & Sakai, R. R. (1995). Serotonin Receptor-Binding in a Colony Model of Chronic Social Stress. Biological Psychiatry, 37(6), 383–393. 10.1016/0006-3223(94)00152-S

McLaren, D. G., Kosmatka, K. J., Kastman, E. K., Bendlin, B. B., & Johnson, S. C. (2010). Rhesus macaque brain morphometry: A methodological comparison of voxel-wise approaches. Methods, 50(3), 157–165. 10.1016/j.ymeth.2009.10.003

Mengod, G., Pompeiano, M., Martinez-Mir, M. I., & Palacios, J. M. (1990). Localization of the mRNA for the 5-HT2 receptor by in situ hybridization histochemistry. Correlation with the distribution of receptor sites. Brain Research, 524(1), 139–143. 10.1016/0006-8993(90)90502-3

Meunier, C. N. J., Amar, M., Lanfumey, L., Hamon, M., & Fossier, P. (2013). 5-HT1A receptors direct the orientation of plasticity in layer 5 pyramidal neurons of the mouse prefrontal cortex. Neuropharmacology, 71, 37–45. 10.1016/j.neuropharm.2013.03.003

Michopoulos, V., Embree, M., Reding, K., Sanchez, M. M., Toufexis, D., Votaw, J. R., Voll, R. J., Goodman, M. M., Rivier, J., Wilson, M. E., & Berga, S. L. (2013). Crh Receptor Antagonism Reverses the Effect of Social Subordination Upon Central Gaba Receptor Binding in Estradiol-Treated Ovariectomized Female Rhesus Monkeys. Neuroscience, 250, 300–308. 10.1016/j.neuroscience.2013.07.002

Michopoulos, V., Perez Diaz, M., Embree, M., Reding, K., Votaw, J. R., Mun, J., Voll, R. J., Goodman, M. M., Wilson, M., Sanchez, M., & Toufexis, D. (2014). Oestradiol alters central 5-HT1A receptor binding potential differences related to psychosocial stress but not differences related to 5-HTTLPR genotype in female rhesus monkeys. J Neuroendocrinol, 26(2), 80–88. 10.1111/jne.12129

Miczek, K. A., Yap, J. J., & Covington, H. E. (2008). Social stress, therapeutics and drug abuse: Preclinical models of escalated and depressed intake. Pharmacology & Therapeutics, 120(2), 102–128. 10.1016/j.pharmthera.2008.07.006

Mocci, G., Jimánez-Sánchez, L., Adell, A., Cortés, R., & Artigas, F. (2014). Expression of 5-HT2A receptors in prefrontal cortex pyramidal neurons projecting to nucleus accumbens. Potential relevance for atypical antipsychotic action. Neuropharmacology, 79, 49–58. 10.1016/j.neuropharm.2013.10.021

Moffett, M. C., Vicentic, A., Kozel, M., Plotsky, P., Francis, D. D., & Kuhar, M. J. (2007). Maternal separation alters drug intake patterns in adulthood in rats. Biochemical Pharmacology, 73(3), 321–330. 10.1016/j.bcp.2006.08.003

Monroy, E., Hernández-Torres, E., & Flores, G. (2010). Maternal separation disrupts dendritic morphology of neurons in prefrontal cortex, hippocampus, and nucleus accumbens in male rat offspring. Journal of Chemical Neuroanatomy, 40(2), 93–101. 10.1016/j.jchemneu.2010.05.005

Morin, E. L., Howell, B. R., Feczko, E., Earl, E., Pincus, M., Reding, K., Kovacs-Balint, Z. A., Meyer, J. S., Styner, M., Fair, D., & Sanchez, M. M. (2020). Developmental outcomes of early adverse care on amygdala functional connectivity in nonhuman primates. Development and Psychopathology, 32(5), 1579–1596. 10.1017/S0954579420001133

Morin, E. L., Howell, B. R., Meyer, J. S., & Sanchez, M. M. (2019). Effects of early maternal care on adolescent attention bias to threat in nonhuman primates. Developmental Cognitive Neuroscience, 38. 10.1016/j.dcn.2019.100643

Morin, E. L., Siebert, E. R., Howell, B. R., Higgins, M., Jovanovic, T., Kazama, A. M., & Sanchez, M. M. (2025). Effects of early maternal care on anxiety and threat learning in adolescent nonhuman primates. Developmental Cognitive Neuroscience, 71. 10.1016/j.dcn.2024.101480

Mukherjee, J., Yang, Z. Y., Brown, T., Lew, R., Wernick, M., Ouyang, X. H., Yasillo, N., Chen, C. T., Mintzer, R., & Cooper, M. (1999). Preliminary assessment of extrastriatal dopamine D-2 receptor binding in the rodent and nonhuman primate brains using the high affinity radioligand,18F-fallypride. Nuclear Medicine and Biology, 26(5), 519–527. 10.1016/S0969-8051(99)00012-8

Muller, C. P., Carey, R. J., Huston, J. P., & De Souza Silva, M. A. (2007). Serotonin and psychostimulant addiction: focus on 5-HT1A-receptors. Prog Neurobiol, 81(3), 133–178. 10.1016/j.pneurobio.2007.01.001

Nader, M. A., Morgan, D., Gage, H. D., Nader, S. H., Calhoun, T. L., Buchheimer, N., Ehrenkaufer, R., & Mach, R. H. (2006). PET imaging of dopamine D2 receptors during chronic cocaine self-administration in monkeys. Nature Neuroscience, 9(8), 1050–1056. 10.1038/nn1737

Nash, J. R., Sargent, P. A., Rabiner, E. A., Hood, S. D., Argyropoulos, S. V., Potokar, J. P., Grasby, P. M., & Nutt, D. J. (2008). Serotonin 5-HT_1A_ receptor binding in people with panic disorder:: positron emission tomography study. British Journal of Psychiatry, 193(3), 229–234. 10.1192/bjp.bp.107.041186

Norberg, M. M., Norton, A. R., Olivier, J., & Zvolensky, M. J. (2010). Social Anxiety, Reasons for Drinking, and College Students. Behavior Therapy, 41(4), 555–566. 10.1016/j.beth.2010.03.002

Öngür, D., & Price, J. L. (2000). The organization of networks within the orbital and medial prefrontal cortex of rats, monkeys and humans. Cerebral Cortex, 10(3), 206–219. 10.1093/cercor/10.3.206

Oswald, L. M., Wand, G. S., Kuwabara, H., Wong, D. F., Zhu, S. J., & Brasic, J. R. (2014). History of childhood adversity is positively associated with ventral striatal dopamine responses to amphetamine. Psychopharmacology, 231(12), 2417–2433. 10.1007/s00213-013-3407-z

Palomero-Gallagher, N., Vogt, B. A., Schleicher, A., Mayberg, H. S., & Zilles, K. (2009). Receptor Architecture of Human Cingulate Cortex: Evaluation of the Four-Region Neurobiological Model. Human Brain Mapping, 30(8), 2336–2355. 10.1002/hbm.20667

Panikratova, Y. R., Vlasova, R. M., Akhutina, T. V., Korneev, A. A., Sinitsyn, V. E., & Pechenkova, E. V. (2020). Functional connectivity of the dorsolateral prefrontal cortex contributes to different components of executive functions. International Journal of Psychophysiology, 151, 70–79. 10.1016/j.ijpsycho.2020.02.013

Pantoja-Urbán, A. H., Richer, S., Mittermaier, A., Giroux, M., Nouel, D., Hernandez, G., & Flores, C. (2024). Gains and Losses: Resilience to Social Defeat Stress in Adolescent Female Mice. Biological Psychiatry, 95(1), 37–47. 10.1016/j.biopsych.2023.06.014

Pazos, A., & Palacios, J. M. (1985). Quantitative autoradiographic mapping of serotonin receptors in the rat brain. I. Serotonin-1 receptors. Brain Research, 346(2), 205–230. 10.1016/0006-8993(85)90856-x

Peña, C. J., Neugut, Y. D., Calarco, C. A., & Champagne, F. A. (2014). Effects of maternal care on the development of midbrain dopamine pathways and reward-directed behavior in female offspring. European Journal of Neuroscience, 39(6), 946–956. 10.1111/ejn.12479

Pentkowski, N. S., Harder, B. G., Brunwasser, S. J., Bastle, R. M., Peartree, N. A., Yanamandra, K., Adams, M. D., Der-Ghazarian, T., & Neisewander, J. L. (2014). Pharmacological Evidence for an Abstinence-Induced Switch in 5-HT1B Receptor Modulation of Cocaine Self-Administration and Cocaine-Seeking Behavior. Acs Chemical Neuroscience, 5(3), 168–176. 10.1021/cn400155t

Perani, D., Garibotto, V., Gorini, A., Moresco, R. M., Henin, M., Panzacchi, A., Matarrese, M., Carpinelli, A., Bellodi, L., & Fazio, F. (2008). In vivo PET study of 5HT(2A) serotonin and D(2) dopamine dysfunction in drug-naive obsessive-compulsive disorder. Neuroimage, 42(1), 306–314. 10.1016/j.neuroimage.2008.04.233

Pilowsky, D. J., Keyes, K. M., & Hasin, D. S. (2009). Adverse Childhood Events and Lifetime Alcohol Dependence. American Journal of Public Health, 99(2), 258–263. 10.2105/Ajph.2008.139006

Pivonello, R., Ferone, D., Lombardi, G., Colao, A., Lamberts, S. W. J., & Hofland, L. J. (2007). Novel insights in dopamine receptor physiology. European Journal of Endocrinology, 156, S13–S21. 10.1530/eje.1.02353

Plaisance, C. J., Ledet, L. I. I. I., Slusher, N. J., Daniel, C. P., Lee, Z. C. Y., Dorius, B., Barrie, S., Parker-Actlis, T. Q., Ahmadzadeh, S., Shekoohi, S., & Kaye, A. D. (2024). The Role of Dopamine in Impulsivity and Substance Abuse: A Narrative Review. Health Psychology Research, 12. 10.52965/001c.125273

Quaedflieg, C. W. E. M., van de Ven, V., Meyer, T., Siep, N., Merckelbach, H., & Smeets, T. (2015). Temporal Dynamics of Stress-Induced Alternations of Intrinsic Amygdala Connectivity and Neuroendocrine Levels. Plos One, 10(5). 10.1371/journal.pone.0124141

Ramboz, S., Oosting, R., Amara, D. A., Kung, H. F., Blier, P., Mendelsohn, M., Mann, J. J., Brunner, D., & Hen, R. (1998). Serotonin receptor 1A knockout: An animal model of anxiety-related disorder. Proceedings of the National Academy of Sciences of the United States of America, 95(24), 14476–14481. 10.1073/pnas.95.24.14476

Rasmussen, S. G. F., Carroll, F. I., Maresch, M. J., Jensen, A. D., Tate, C. G., & Gether, U. (2001). Biophysical characterization of the cocaine binding pocket in the serotonin transporter using a fluorescent cocaine analogue as a molecular reporter. Journal of Biological Chemistry, 276(7), 4717–4723. 10.1074/jbc.M008067200

Rentesi, G., Antoniou, K., Marselos, M., Syrrou, M., Papadopoulou-Daifoti, Z., & Konstandi, M. (2013). Early maternal deprivation-induced modifications in the neurobiological, neurochemical and behavioral profile of adult rats. Behavioural Brain Research, 244, 29–37. 10.1016/j.bbr.2013.01.040

Reynolds, L. M., & Flores, C. (2021). Mesocorticolimbic Dopamine Pathways Across Adolescence: Diversity in Development. Frontiers in Neural Circuits, 15. 10.3389/fncir.2021.735625

Richardson, N. R., & Roberts, D. C. S. (1996). Progressive ratio schedules in drug self-administration studies in rats: A method to evaluate reinforcing efficacy. Journal of Neuroscience Methods, 66(1), 1–11. 10.1016/0165-0270(95)00153-0

Rodrigues, S. M., LeDoux, J. E., & Sapolsky, R. M. (2009). The Influence of Stress Hormones on Fear Circuitry. Annual Review of Neuroscience, 32, 289–313. 10.1146/annurev.neuro.051508.135620

Sanchez, E. O., Bavley, C. C., Deutschmann, A. U., Carpenter, R., Peterson, D. R., Karbalaei, R., Flowers, J., Rogers, C. M., Langrehr, M. G., Ardekani, C. S., Famularo, S. T., Bongiovanni, A. R., Knouse, M. C., Floresco, S. B., Briand, L. A., Wimmer, M. E., & Bangasser, D. A. (2022). Early life adversity promotes resilience to opioid addiction-related phenotypes in male rats and sex-specific transcriptional changes (vol 118, e2020173118, 2021). Proceedings of the National Academy of Sciences of the United States of America, 119(17). 10.1073/pnas.2204210119

Sanchez, M. M. (2006). The impact of early adverse care on HPA axis development: nonhuman primate models. Horm Behav, 50(4), 623–631. 10.1016/j.yhbeh.2006.06.012

Sanchez, M. M., Ladd, C. O., & Plotsky, P. M. (2001). Early adverse experience as a developmental risk factor for later psychopathology: Evidence from rodent and primate models. Development and Psychopathology, 13(3), 419–449. 10.1017/S0954579401003029

Savitz, J. B., & Drevets, W. C. (2013). Neuroreceptor imaging in depression. Neurobiology of Disease, 52, 49–65. 10.1016/j.nbd.2012.06.001

Schenk, S., & Highgate, Q. (2021). Methylenedioxymethamphetamine (MDMA): Serotonergic and dopaminergic mechanisms related to its use and misuse. Journal of Neurochemistry, 157(5), 1714–1724. 10.1111/jnc.15348

Schwartz, E. K., Wolkowicz, N. R., De Aquino, J. P., MacLean, R. R., & Sofuoglu, M. (2022). Cocaine Use Disorder (CUD): Current Clinical Perspectives. Substance Abuse and Rehabilitation, 13, 25–46. 10.2147/Sar.S337338

Sharp, T., & Barnes, N. M. (2020). Central 5-HT receptors and their function; present and future. Neuropharmacology, 177. 10.1016/j.neuropharm.2020.108155

Shi, Y. D., Budin, F., Yapuncich, E., Rumple, A., Young, J. T., Payne, C., Zhang, X. D., Hu, X. P., Godfrey, J., Howell, B., Sanchez, M. M., & Styner, M. A. (2017). UNC-Emory Infant Atlases for Macaque Brain Image Analysis: Postnatal Brain Development through 12 Months. Frontiers in Neuroscience, 10. 10.3389/fnins.2016.00617

Simpson, T. L., & Miller, W. R. (2002). Concomitance between childhood sexual and physical abuse and substance use problems - A review. Clinical Psychology Review, 22(1), 27–77. 10.1016/S0272-7358(00)00088-X

Sinha, R. (2008). Chronic Stress, Drug Use, and Vulnerability to Addiction. Addiction Reviews 2008, 1141, 105–130. 10.1196/annals.1441.030

Sinha, R., Lacadie, C. M., Constable, R. T., & Seo, D. (2016). Dynamic neural activity during stress signals resilient coping. Proceedings of the National Academy of Sciences of the United States of America, 113(31), 8837–8842. 10.1073/pnas.1600965113

Smith, K. E., & Pollak, S. D. (2020). Early life stress and development: potential mechanisms for adverse outcomes. Journal of Neurodevelopmental Disorders, 12(1). 10.1186/s11689-020-09337-y

Solinas, M., Belujon, P., Fernagut, P. O., Jaber, M., & Thiriet, N. (2019). Dopamine and addiction: what have we learned from 40 years of research. Journal of Neural Transmission, 126(4), 481–516. 10.1007/s00702-018-1957-2

Spinelli, S., Chefer, S., Carson, R. E., Jagoda, E., Lang, L. X., Heilig, M., Barr, C. S., Suomi, S. J., Higley, J. D., & Stein, E. A. (2010). Effects of Early-Life Stress on Serotonin_1A_ Receptors in Juvenile Rhesus Monkeys Measured by Positron Emission Tomography. Biological Psychiatry, 67(12), 1146–1153. 10.1016/j.biopsych.2009.12.030

Stockmeier, C. A. (2003). Involvement of serotonin in depression: evidence from postmortem and imaging studies of serotonin receptors and the serotonin transporter. Journal of Psychiatric Research, 37(5), 357–373. 10.1016/S0022-3956(03)00050-5

Stuber, G. D., Sparta, D. R., Stamatakis, A. M., van Leeuwen, W. A., Hardjoprajitno, J. E., Cho, S., Tye, K. M., Kempadoo, K. A., Zhang, F., Deisseroth, K., & Bonci, A. (2011). Excitatory transmission from the amygdala to nucleus accumbens facilitates reward seeking. Nature, 475(7356), 377–U129. 10.1038/nature10194

Sun, S., Yu, H. B., Yu, R. J., & Wang, S. (2023). Functional connectivity between the amygdala and prefrontal cortex underlies processing of emotion ambiguity. Translational Psychiatry, 13(1). 10.1038/s41398-023-02625-w

Teicher, M. H., Andersen, S. L., Polcari, A., Anderson, C. M., Navalta, C. P., & Kim, D. M. (2003). The neurobiological consequences of early stress and childhood maltreatment. Neurosci Biobehav Rev, 27(1-2), 33–44. 10.1016/s0149-7634(03)00007-1

Teicher, M. H., & Samson, J. A. (2016). Annual Research Review: Enduring neurobiological effects of childhood abuse and neglect. Journal of Child Psychology and Psychiatry, 57(3), 241–266. 10.1111/jcpp.12507

Turkheimer, F. E., Selvaraj, S., Hinz, R., Murthy, V., Bhagwagar, Z., Grasby, P., Howes, O., Rosso, L., & Bose, S. K. (2012). Quantification of ligand PET studies using a reference region with a displaceable fraction: application to occupancy studies with [^11^C]-DASB as an example. Journal of Cerebral Blood Flow and Metabolism, 32(1), 70–80. 10.1038/jcbfm.2011.108

Varnäs, K., Halldin, C., & Hall, H. (2004). Autoradiographic distribution of serotonin transporters and receptor subtypes in human brain. Human Brain Mapping, 22(3), 246–260. 10.1002/hbm.20035

Vassilev, P., Pantoja-Urban, A. H., Giroux, M., Nouel, D., Hernandez, G., Orsini, T., & Flores, C. (2021). Unique Effects of Social Defeat Stress in Adolescent Male Mice on the Netrin-1/DCC Pathway, Prefrontal Cortex Dopamine and Cognition. Eneuro, 8(2). 10.1523/Eneuro.0045-21.2021

Vazquez, V., Penit-Soria, J., Durand, C., Besson, M. J., Giros, B., & Daugé, V. (2006). Brief early handling increases morphine dependence in adult rats. Behavioural Brain Research, 170(2), 211–218. 10.1016/j.bbr.2006.02.022

Vázquez-Borsetti, P., Cortés, R., & Artigas, F. (2009). Pyramidal Neurons in Rat Prefrontal Cortex Projecting to Ventral Tegmental Area and Dorsal Raphe Nucleus Express 5-HT2A Receptors. Cerebral Cortex, 19(7), 1678–1686. 10.1093/cercor/bhn204

Verma, V. (2015). Classic Studies on the Interaction of Cocaine and the Dopamine Transporter. Clinical Psychopharmacology and Neuroscience, 13(3), 227–238. 10.9758/cpn.2015.13.3.227

Volkow, N. D., & Boyle, M. (2018). Neuroscience of Addiction: Relevance to Prevention and Treatment. American Journal of Psychiatry, 175(8), 729–740. 10.1176/appi.ajp.2018.17101174

Volkow, N. D., Michaelides, M., & Baler, R. (2019). The Neuroscience of Drug Reward and Addiction. Physiological Reviews, 99(4), 2115–2140. 10.1152/physrev.00014.2018

Wakeford, A. G. P., Kochoian, B., Siebert, E. R., Katznelson, S., Morin, E. L., Howell, B. R., McCormack, K. M., Nader, M. A., Sanchez, M. M., & Howell, L. L. (2020). Effects of early life stress on cocaine intake in male and female rhesus macaques. Psychopharmacology. 10.1007/s00213-020-05637-2

Wakeford, A. G. P., Morin, E. L., Bramlett, S. N., Howell, B. R., McCormack, K. M., Meyer, J. S., Nader, M. A., Sanchez, M. M., & Howell, L. L. (2019). Effects of early life stress on cocaine self-administration in post-pubertal male and female rhesus macaques. Psychopharmacology (Berl), 236(9), 2785–2796. 10.1007/s00213-019-05254-8

Wakeford, A. G. P., Morin, E. L., Bramlett, S. N., Howell, L. L., & Sanchez, M. M. (2018). A review of nonhuman primate models of early life stress and adolescent drug abuse. Neurobiol Stress, 9, 188–198. 10.1016/j.ynstr.2018.09.005

Wakeford, A. G. P., Nye, J. A., Morin, E. L., Mun, J., Meyer, J. S., Goodman, M., Howell, L. L., & Sanchez, M. M. (2024). Alterations in adolescent brain serotonin (5HT)_1A_, 5HT_2A_, and dopamine (D)_2_ receptor systems in a nonhuman primate model of early life adversity. Neuropsychopharmacology, 49(8), 1227–1235. 10.1038/s41386-023-01784-0

Wand, G. S., Oswald, L. M., McCaul, M. E., Wong, D. F., Johnson, E., Zhou, Y., Kuwabara, H., & Kumar, A. (2007). Association of amphetamine-induced striatal dopamine release and cortisol responses to psychological stress. Neuropsychopharmacology, 32(11), 2310–2320. 10.1038/sj.npp.1301373

Watanabe, Y., Sakai, R. R., McEwen, B. S., & Mendelson, S. (1993). Stress and antidepressant effects on hippocampal and cortical 5-HT1A and 5-HT2 receptors and transport sites for serotonin. Brain Research, 615(1), 87–94. 10.1016/0006-8993(93)91117-b

Widom, C. S. (1999). Posttraumatic stress disorder in abused and neglected children grown up. American Journal of Psychiatry, 156(8), 1223–1229.

Woods, R. P., Cherry, S. R., & Mazziotta, J. C. (1992). Rapid Automated Algorithm for Aligning and Reslicing Pet Images. Journal of Computer Assisted Tomography, 16(4), 620–633. 10.1097/00004728-199207000-00024

Xu, P., Chen, A., Li, Y. P., Xing, X. Z., & Lu, H. (2019). Medial prefrontal cortex in neurological diseases. Physiological Genomics, 51(9), 432–442. 10.1152/physiolgenomics.00006.2019

Xu, Y., Lin, Y. J., Yu, M., & Zhou, K. K. (2024). The nucleus accumbens in reward and aversion processing: insights and implications. Frontiers in Behavioral Neuroscience, 18. 10.3389/fnbeh.2024.1420028

Yalcin, I., Barthas, F., & Barrot, M. (2014). Emotional consequences of neuropathic pain: Insight from preclinical studies. Neuroscience and Biobehavioral Reviews, 47, 154–164. 10.1016/j.neubiorev.2014.08.002

Yan, C. G., Rincón-Cortés, M., Raineki, C., Sarro, E., Colcombe, S., Guilfoyle, D. N., Yang, Z., Gerum, S., Biswal, B. B., Milham, M. P., Sullivan, R. M., & Castellanos, F. X. (2017). Aberrant development of intrinsic brain activity in a rat model of caregiver maltreatment of offspring. Translational Psychiatry, 7. 10.1038/tp.2016.276

Yang, Y. F., Tai, Y. C., Siegel, S., Newport, D. F., Bai, B., Li, Q. Z., Leahy, R. M., & Cherry, S. R. (2004). Optimization and performance evaluation of the microPET II scanner for small-animal imaging. Physics in Medicine and Biology, 49(12), 2527–2545. 10.1088/0031-9155/49/12/005

Young, R. F., & Tandon, M. (2025). Child Maltreatment A Review on Prevention, Intervention, and Impact. Child and Adolescent Psychiatric Clinics of North America, 34(2), 311–323. 10.1016/j.chc.2024.08.006

Zhu, H., Wang, S., Qu, L., & Shen, D. (2021). Chapter 10 - Hippocampus segmentation in MR images: Multiatlas methods and deep learning methods. In A. A. Moustafa (Ed.), Big Data in Psychiatry #x0026; Neurology (pp. 181–215). Academic Press. 10.1016/B978-0-12-822884-5.00019-2

