## Supplemental Material for "Long-term Effects of Early Life Adversity On Brain Dopamine and Serotonin Receptor Systems Involved In Cocaine Reinforcement In Adult Macaques: a Positron Emission Tomography Study"

### **Supplemental Materials**

**Supplemental Tables:**

**Suppl. Table 1. Spearman’s Rank Correlations Between ROIs PET Binding Potentials (BP) at Baseline (pre-cocaine(COC) self-administration (SA)) for 5HT_1A_ receptor BP.** ROIs: ACC, Amygdala, Caudate, Hippocampus, NAcc, OFC, Putamen, SGC, dlPFC, mPFC, vlPFC, and vmPFC.

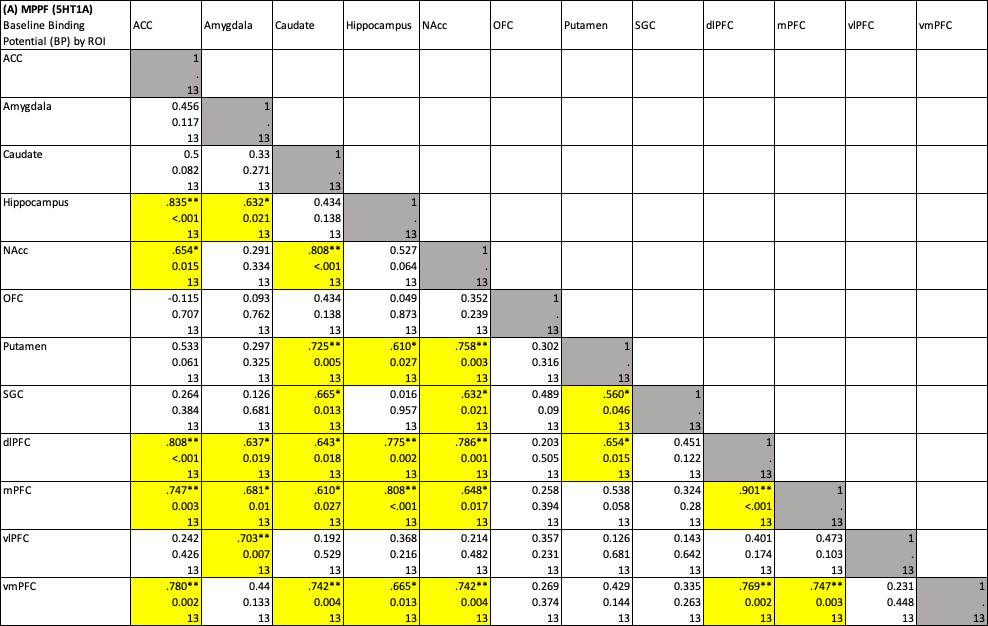

Cells represent rho value, p value (*: *p*<.05, **: *p*<.01), and sample size (n).

**Suppl. Table 2. Spearman’s Rank Correlations Between ROIs PET Binding Potentials (BP) at Baseline (pre-cocaine self-administration) for 5HT_2A_ receptor BP.** ROIs: ACC, Amygdala, Caudate, Hippocampus, NAcc, OFC, Putamen, SGC, dlPFC, mPFC, vlPFC, and vmPFC.

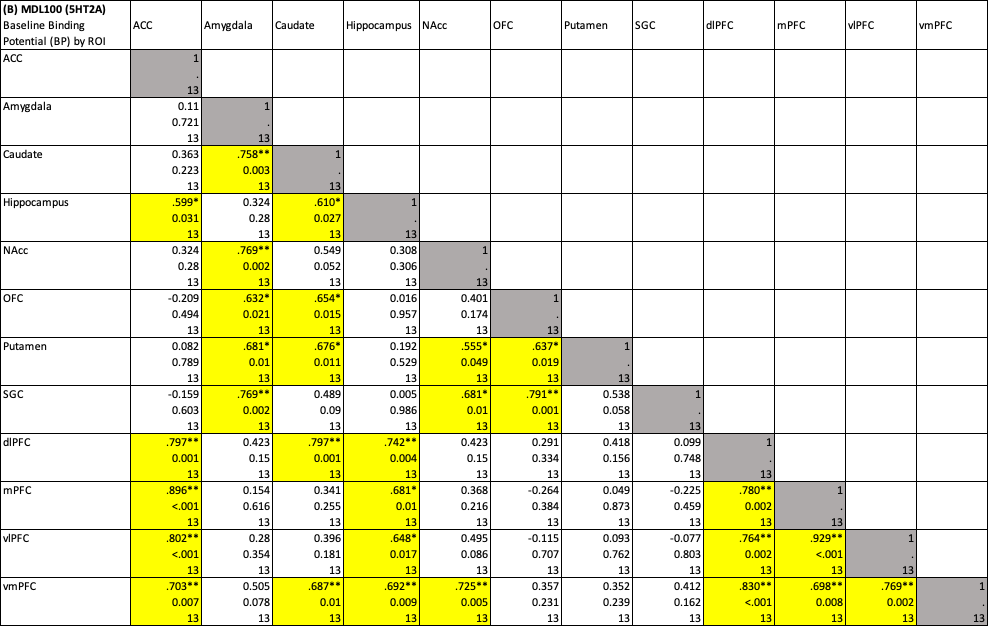

Cells represent rho value, p value (*: *p*<.05, **: *p*<.01), and sample size (n).

**Suppl. Table 3. Spearman’s Rank Correlations Between ROIs PET Binding Potentials (BP) at Baseline (pre-cocaine self-administration) for D_2_/D_3_ receptor BP.** ROIs: ACC, Amygdala, Caudate, Hippocampus, NAcc, OFC, Putamen, SGC, dlPFC, mPFC, vlPFC, and vmPFC.

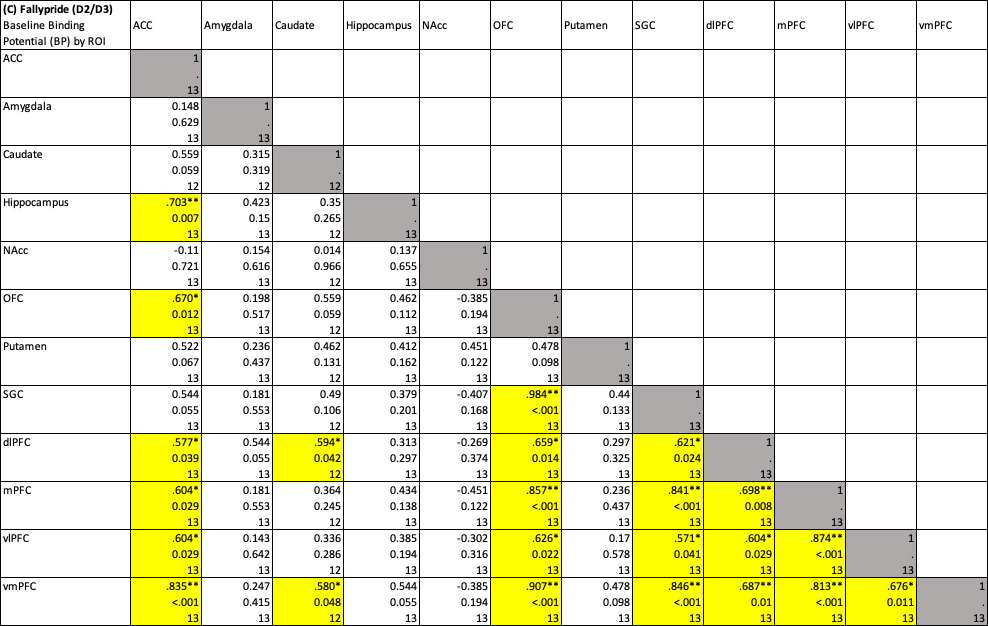

Cells represent rho value, p value (*: *p*<.05, **: *p*<.01), and sample size (n).

### **Supplemental Figures:**

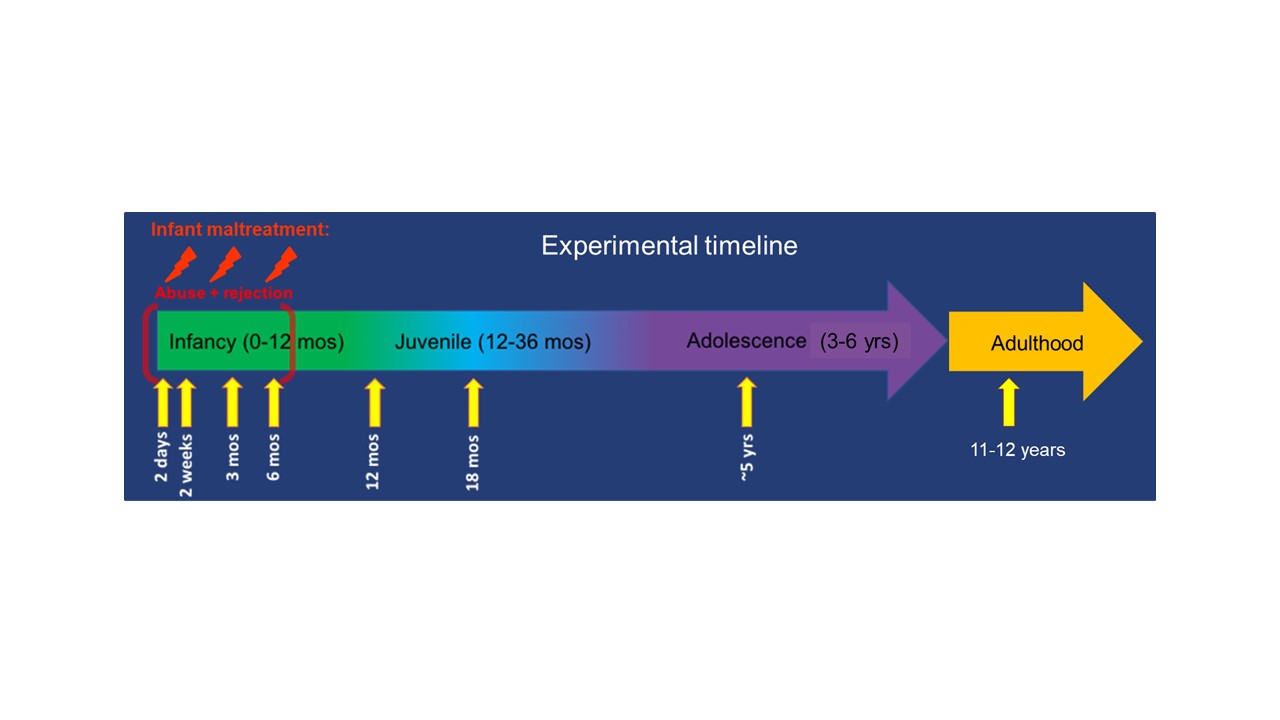

**Suppl. Figure 1. Experimental Timeline.** At birth, animals were randomly assigned to experimental group (Control, Maltreated) based on their foster mom’ maternal care history. Maternal care data received from foster mom was collected for 3 months after birth to confirm MALT/ELS experience and rates of abuse and rejection, in contrast to nurturing maternal behaviors. Infants were studied longitudinally from infancy through adulthood (current study).

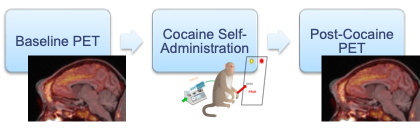

**Suppl. Figure 2. PET scan progression*.*** These PET scans were acquired at (1) baseline (pre-COC SA) in all 22 adult animals, and (2) again, post-COC SA, right after completion of post-COC SA FR and PR studies, and once each individual had reached a cumulative COC intake of ≥100mgCOC/kg BW.

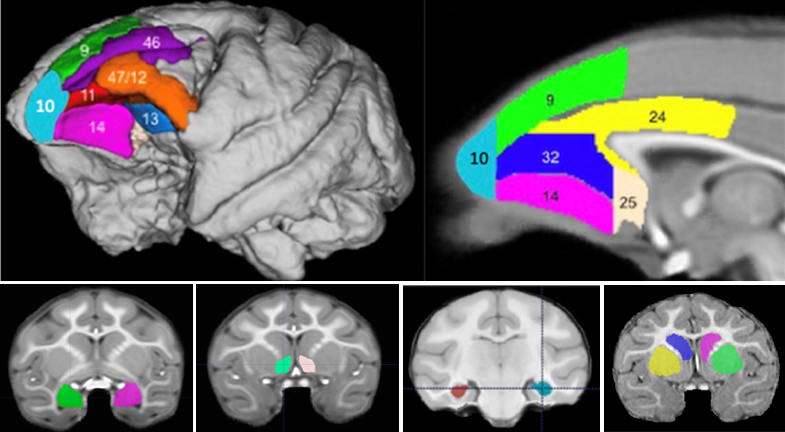

**Suppl. Figure 3. *Regions of interest (ROIs) displayed in the rhesus brain MRI atlas.*** (Top-left) Lateral surface view of prefrontal cortex (PFC) subregions in the rhesus brain MRI atlas image. (Top-right) Sagittal view of additional PFC ROIs in the rhesus brain MRI atlas image. Those PFC areas were merged into the following PFC subregions ROIs for PET BP analyses: anterior cingulate cortex -ACC: area 24-, subgenual cingulate cortex -SGC: area 25- medial PFC -mPFC: areas 10 and 32-, ventromedial PFC -vmPFC: area 14-, orbitofrontal cortex -OFC: areas 11 and 13-, dlPFC -areas 9 and 46-, and ventrolateral PFC -vlPFC: area 12/47-. Coronal views of the rest of the ROIs in the rhesus brain MRI atlas image (Bottom Left to Right): amygdala, nucleus Accumbens -NAcc-, hippocampus, caudate (left: blue; right: pink)/putamen(left: yellow/right: green). Figures of PFC subregions reproduced from (Kovacs-Balint et al., 2023) with minor changes (left/right images swapped and label added for area 10).
